# Multi-omics reveals lipid metabolism, mitochondrial and extracellular matrix dysregulation of diabetic cardiomyopathy in the human heart

**DOI:** 10.1101/2025.08.21.671651

**Authors:** Qiuhan Lu, Shulin Tang, Yuwen Wu, Liang Chen, Mintong Liang, Sijia Fang, Jiaqi Chen, Pengju Wen, Leigang Jin, Jianshe Yu, Feng Jiao, Yueheng Wu, Guozhi Jiang

## Abstract

Diabetic cardiomyopathy (DbCM) is a condition characterized by myocardial dysfunction in diabetes. Although the clinical recognition of DbCM is increasing, its underlying molecular mechanisms remain poorly understood. We aimed to conduct a comprehensive multi-omics analysis of heart tissue from cardiomyopathy patients (both with and without diabetes),alongside tissue from healthy donors, by integrating transcriptomic, 4D-DIA proteomic, and full-spectrum widely targeted metabolomic data. Differential analyses revealed a significant metabolic pattern in patients with DbCM, characterized by increasing utilization of triglyceride-derived fatty acids. Correlation analyses highlighted impaired BNIP3L-mediated mitophagy as a potential contributor to the disruption of fatty acid metabolism and mitochondrial dysfunction in DbCM. Protein interaction and fibrotic trichrome analyses showed that patients with DbCM had impaired cardiac matrix repair compared to patients with cardiomyopathy without diabetes. These findings provide a system-level understanding of DbCM and reveal molecular signatures driving its progression, offering insights for targeted therapeutic strategies against diabetic heart disease.

## Introduction

The hyperglycemic environment and insulin resistance characteristics of type 2 diabetes (T2D) detrimentally affect the myocardial structure and function, either independently or in conjunction with cardiovascular disease [1, 2]. The escalating global prevalence of T2D is particularly concerning, given that individuals with T2D exhibit a two- to five-fold higher risk of developing heart failure (HF) than non-diabetic individuals [3, 4]. Chronic inflammation and oxidative stress, common features of T2D, further predispose patients to microcirculatory alterations and cardiovascular disease [1, 5]. Among cardiovascular complications, diabetic cardiomyopathy (DbCM), recently defined as systolic and/or diastolic myocardial dysfunction occurring in the context of diabetes, affects at least 12% of patients with T2D [6, 7, 8]. However, the insidious onset and progression of DbCM often delay diagnoses and interventions, exacerbating the disease burden [5, 9, 10].

The prevention and management of cardiac dysfunction in T2D remains a major clinical challenge, fueling the emergence of a new subspecialty—diabetocardiology [2, 11]. Despite the growing recognition of DbCM, the molecular mechanisms linking diabetes to myocardial injury remain poorly understood, largely because of the limited availability of human heart tissue samples. Recent studies have addressed these issues. Hahn et al. integrated transcriptomic and metabolomic analyses in a cohort of patients with HF, revealing widespread impairments in myocardial metabolic substrate utilization, particularly among patients with obesity and heart failure with preserved ejection fraction [12, 13]. Furthermore, Shu et al. employed targeted metabolomics and transcriptomics to identify the myocardial accumulation of long-chain acyl-coenzyme A as a metabolic hallmark of DbCM [14]. However, despite these advances, the multi-omics landscape of DbCM in the human heart remains largely unexplored.

Rodent models are easy to establish but often fail to capture the complexity of human cardiac physiology and pathology, limiting their translational relevance. Blood-based omics enables convenient, longitudinal sampling but lacks structural resolution. Given the multifactorial nature of DbCM, integrating multi-omics data is essential for a comprehensive understanding of disease mechanisms, molecular interactions, and temporal progression.

To address this knowledge gap, in this study we conducted a multi-omics investigation using left ventricular tissues from Chinese patients with cardiomyopathy (both with and without diabetes) and healthy donors. Our study aimed to comprehensively delineate the pathophysiological alterations associated with DbCM across transcriptomic, 4D-DIA proteomic, and full-spectrum widely targeted metabolomic dimensions. We identified distinct molecular signatures of DbCM characterized by dysregulated lipid metabolism, mitochondrial dysfunction, and extracellular matrix remodeling. These findings provide valuable insights into the mechanisms driving cardiac dysfunction in diabetes and contribute to a deeper mechanistic understanding of the pathophysiology of DbCM in the human heart.

## Results

### Clinical characteristics of patients

Left ventricular tissue samples were collected from 22 patients (11 with DbCM and 11 matched cardiomyopathy patients without diabetes; 81.8% males) and four healthy donors. The baseline characteristics of patients with DbCM and those with cardiomyopathy without diabetes are presented in Table 1. The mean ages of patients in the DbCM and cardiomyopathy without diabetes groups were 57.2±10.2 years and 56.2±8.9 years, respectively. No significant differences were observed in cardiovascular conditions between the two groups. The detailed clinical information of the individuals is provided in Supplementary Table 1.

**Table 1.**
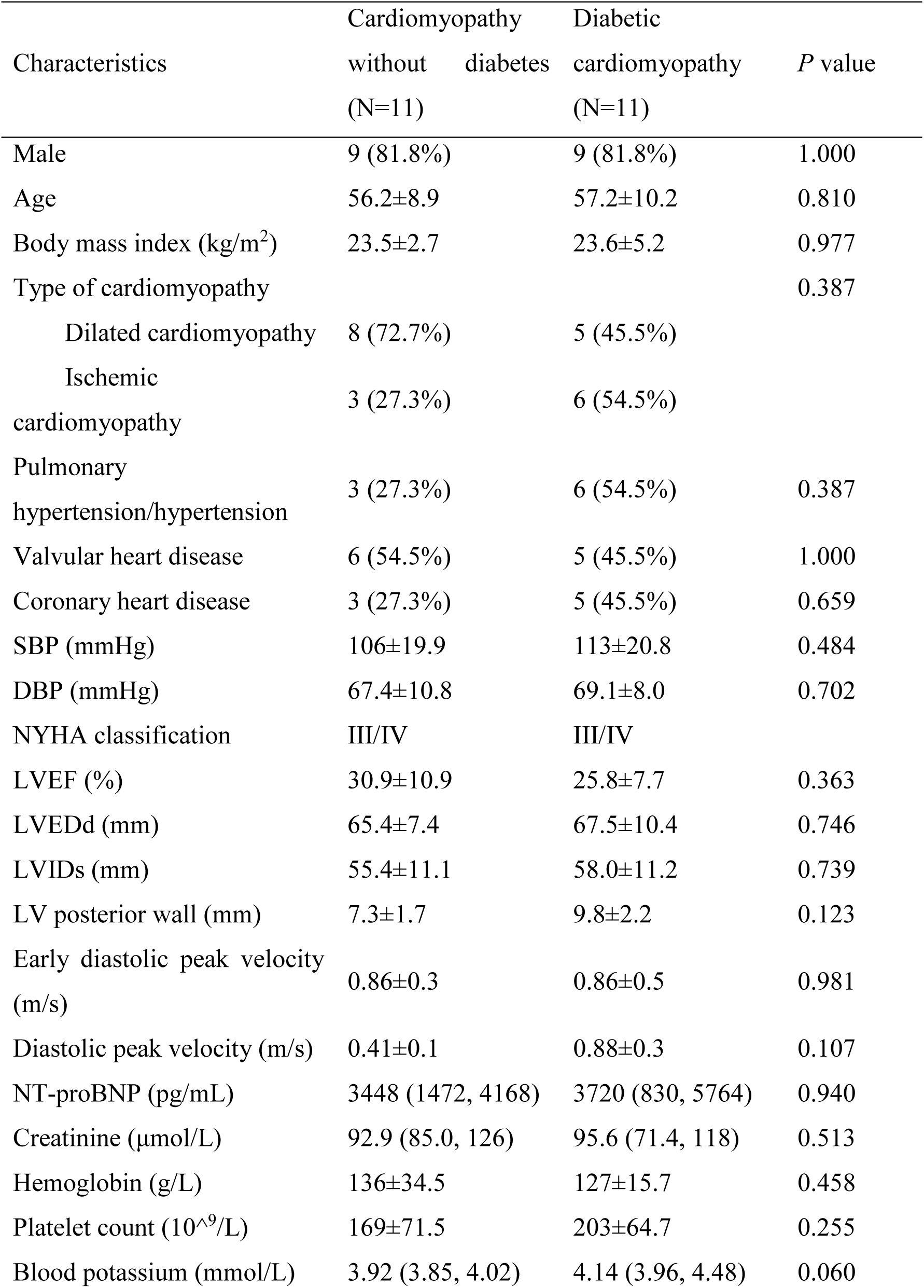
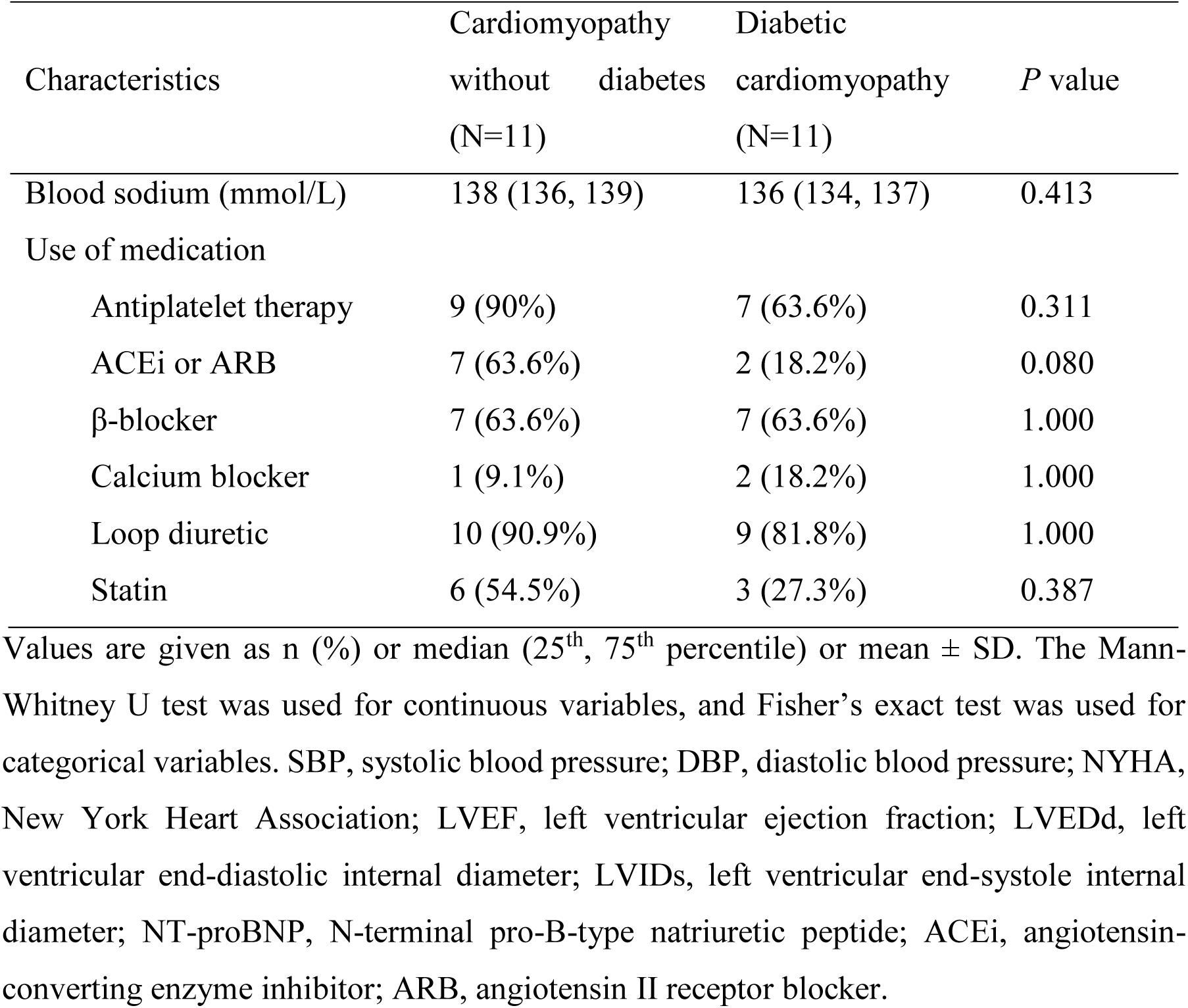
Baseline characteristics of patients.

After quality control and filtration, 15,788 genes, 6,459 proteins, and 4,116 metabolites were retained for multi-omics analysis to characterize the molecular composition of the human heart tissues. However, one sample from the cardiomyopathy without diabetes group was identified as an outlier in the principal component analysis and was subsequently excluded (Supplementary Fig. 1). An overview of the workflow is presented in Fig. 1. Partial least squares discriminant analysis (PLS-DA) revealed distinct multi-omics profiles between the groups (Fig. 2A-C, Source Data 1).

**Fig. 1.**
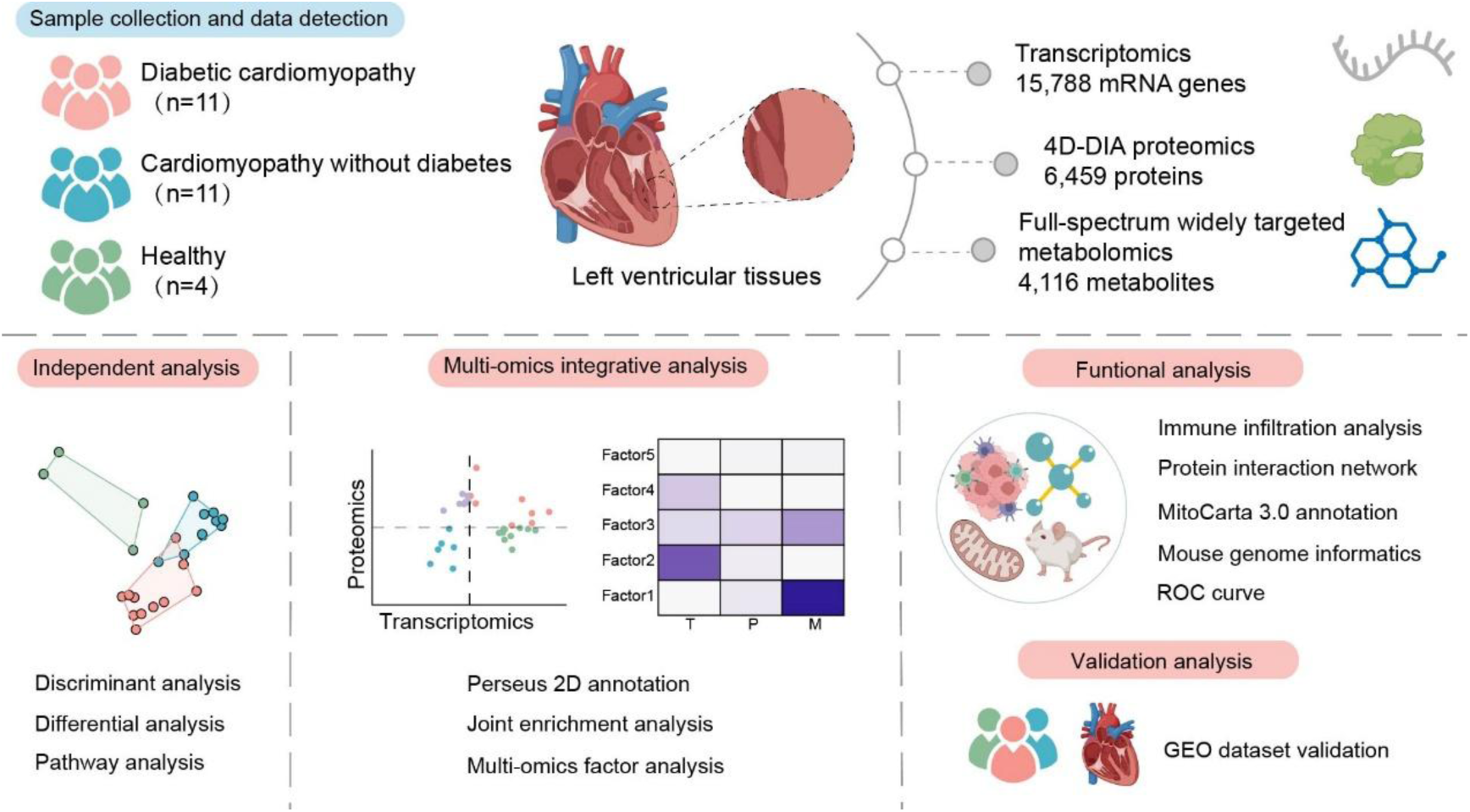
Overview of the multi-omics analysis. The analysis workflow comprised four key stages: construction of the sample collection and multi-omics data acquisition, independent analysis of each omics dataset, integrative multi-omics analysis, and validation using independent cohorts and experiments.

**Fig. 2.**
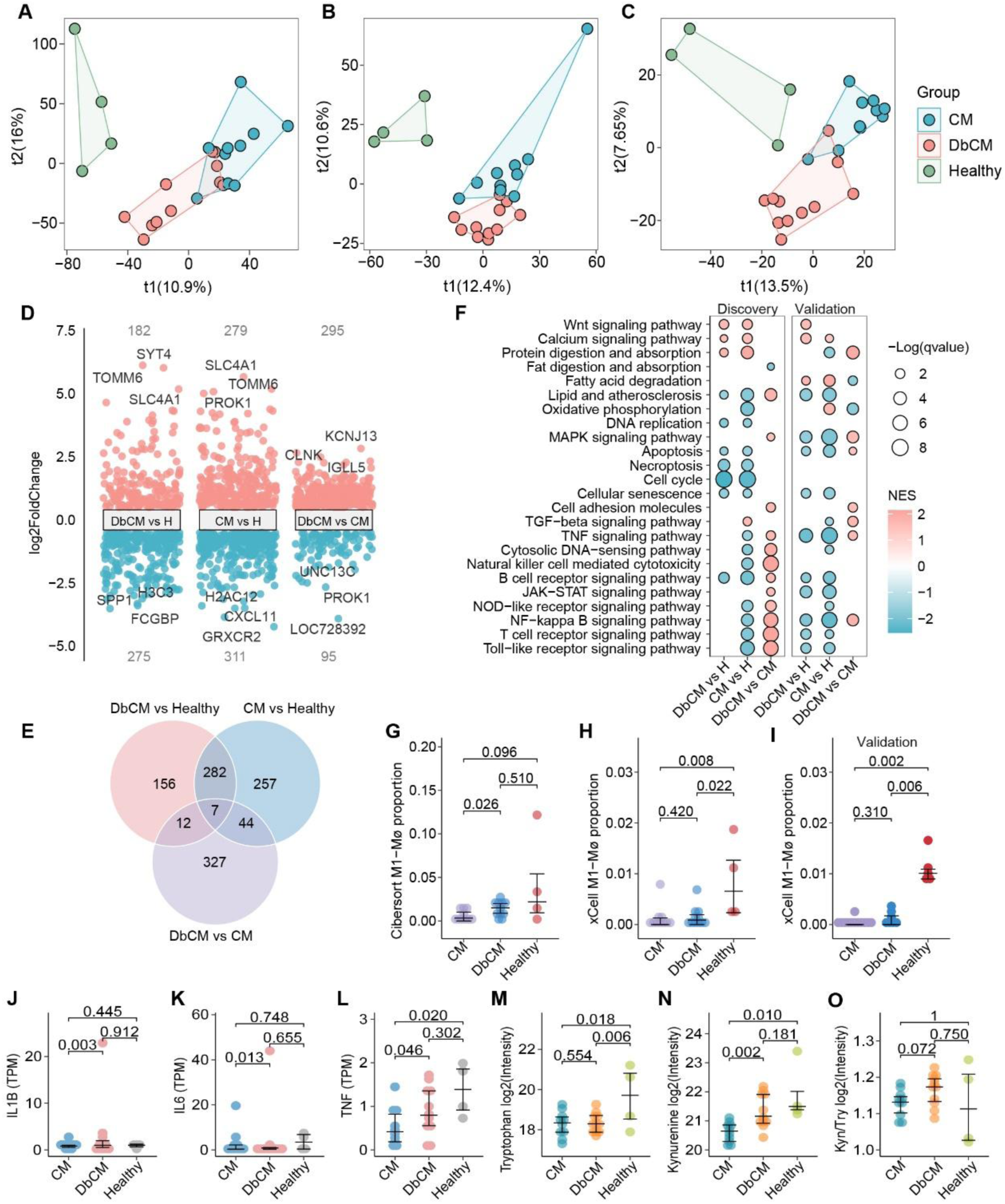
Diabetes activated inflammation and immune pathways in the heart of DbCM patients. (**A-C**) PLS-DA plots of transcriptome, proteome, and metabolome, respectively, demonstrating distinct multi-omics profiles between groups. (**D**) Multi-volcano plot of DEGs across multiple comparison groups. The top three genes with the highest absolute log_2_FoldChange were labeled, and total deg counts were displayed in grey. (**E**) Venn diagram summarizing overlapping DEGs. (**F**) GSEA of transcriptome data from the KEGG database revealed downregulated immune and inflammatory regulation in both disease groups compared to healthy controls. Diabetes activated immune and inflammatory regulation in the hearts of DbCM patients compared to cardiomyopathy patients without diabetes, with external validation supporting these findings. Dot size represents −log(p-value) and color indicates normalized enrichment scores (NES). (**G, H**) Cibersort and xCell analysis showed lower proportions of M1 macrophages (Mø) in both disease groups compared to the healthy group, with *P*-values assessed by the Mann-Whitney U test. The horizontal line represents the median and error range. (**i**) xCell validation analysis confirmed the alterations for M1 macrophages. (**J-L**) TPM values of IL1B, IL6, TNF in cardiomyopathy patients without diabetes, DbCM patients and healthy controls. *P*-values for differential analysis were shown. (**M-O**) Relative intensities of tryptophan, kynurenine, and their ratio.

### Transcriptome landscape

Compared with the healthy group, 457 and 590 genes were dysregulated in the DbCM and cardiomyopathy without diabetes groups, respectively, whereas 390 differentially expressed genes (DEGs) were identified between the DbCM and cardiomyopathy without diabetes groups (Fig. 2D, Source Data 2). Gene set enrichment analysis (GSEA) based on the Kyoto Encyclopedia of Genes and Genomes (KEGG) database revealed activation of the Wnt signaling pathway and protein digestion and absorption in both disease groups compared to healthy controls, whereas pathways associated with immune regulation and inflammation were downregulated (Fig. 2F, Source Data 3). These findings suggest a shared dysregulation of biological processes in patients with DbCM and cardiomyopathy without diabetes (Fig. 2E-F). Further comparison between DbCM and cardiomyopathy without diabetes revealed the activation of pathways related to the immune response, inflammation, and lipid metabolism in the DbCM group (Fig. 2F). CIBERSORT analysis demonstrated a higher proportion of M1 macrophages in patients with DbCM than in those with cardiomyopathy without diabetes, with both disease groups showing lower levels of M1 macrophages than those in the healthy group (Fig. 2G, Source Data 4). The xCell analysis confirmed a similar trend in the estimated proportion of M1 macrophages. (Fig. 2H).

External validation using data from Greco et al. [15] further supported the observation that immune response and inflammation pathways were downregulated in both patient groups, while diabetes was associated with the activation of transforming growth factor (TGF)-β signaling, tumor necrosis factor (TNF) signaling, and nuclear factor-κB signaling pathways in the hearts of patients with DbCM compared to those with cardiomyopathy without diabetes (Fig. 2F). The xCell analysis of the validation data showed a consistent trend in the proportion of M1 macrophages (Fig. 2I).

### Diabetes activated inflammation and immune pathways in the heart of DbCM patients compared to cardiomyopathy without diabetes

While overactivation of inflammation is a critical pathogenic factor in cardiomyopathy [1], both DbCM and cardiomyopathy without diabetes exhibit suppressed immune and inflammatory regulation compared with healthy controls. However, diabetes notably activates inflammation-related pathways in patients compared to patients with cardiomyopathy without diabetes, indicating that it plays a proinflammatory role in the pathophysiology of cardiomyopathy. After adjusting for age, sex, and type of cardiomyopathy, interleukin 1 beta (IL1B) (log_2_FC=1.91, *P*=0.003), interleukin 6 (IL6) (log_2_FC=1.87, *P*=0.013), and TNF (log_2_FC=1.06, *P*=0.046) were significantly upregulated in the transcriptomes of patients with DbCM compared with cardiomyopathy patients without diabetes (Fig. 2J-L). Tryptophan catabolism via the kynurenine pathway has been implicated in chronic inflammatory diseases and may directly influence systemic immunity [16]. Here, tryptophan levels were significantly reduced in patients with DbCM (log_2_FC=-1.21, *P*=0.006) and cardiomyopathy patients without diabetes (log_2_FC=-1.04, *P*=0.018) compared to healthy controls, although no significant differences were observed between the disease groups (Fig. 2M). Furthermore, kynurenine levels were elevated in patients with DbCM compared to those with cardiomyopathy without diabetes (log_2_FC=0.75, *P*=0.002) (Fig. 2N). The tryptophan-to-kynurenine ratio was higher in patients with DbCM (Fig. 2O).

### Proteome profiling

Transcript expression showed a modest correlation with protein levels (r=0.334, *P*<0.001), emphasizing the necessity for proteomic analysis. Analysis of differentially expressed proteins (DEPs) revealed significant proteomic alterations in both the DbCM (236 proteins) and cardiomyopathy without diabetes (273 proteins) groups compared to healthy controls (Fig. 3A-B, Source Data 2). Furthermore, 176 proteins were differentially expressed between the DbCM and cardiomyopathy without diabetes groups after adjusting for age, sex, and cardiomyopathy (Fig. 3A-B). The upregulated DEPs in both the DbCM and cardiomyopathy without diabetes groups were involved in the extracellular matrix (ECM) and fiber organization, as well as in the activation of matrix metalloproteinases (MMPs), when compared to healthy controls (Fig. 3C, Source Data 5). In contrast, DEPs distinguishing DbCM from cardiomyopathy without diabetes showed enrichment in pathways related to acylglycerol metabolism and lipid catabolism, while lipid biosynthetic processes were downregulated (Fig. 3D), indicating marked disruptions in lipid metabolic regulation. Additionally, STEAP3 metalloreductase was downregulated in patients with DbCM relative to healthy controls, with substantial reductions in both mRNA (log_2_FC=-0.87, *P*=0.003) and protein levels (log_2_FC=-0.51, *P*<0.001) (Fig. 3E-F).

**Fig. 3.**
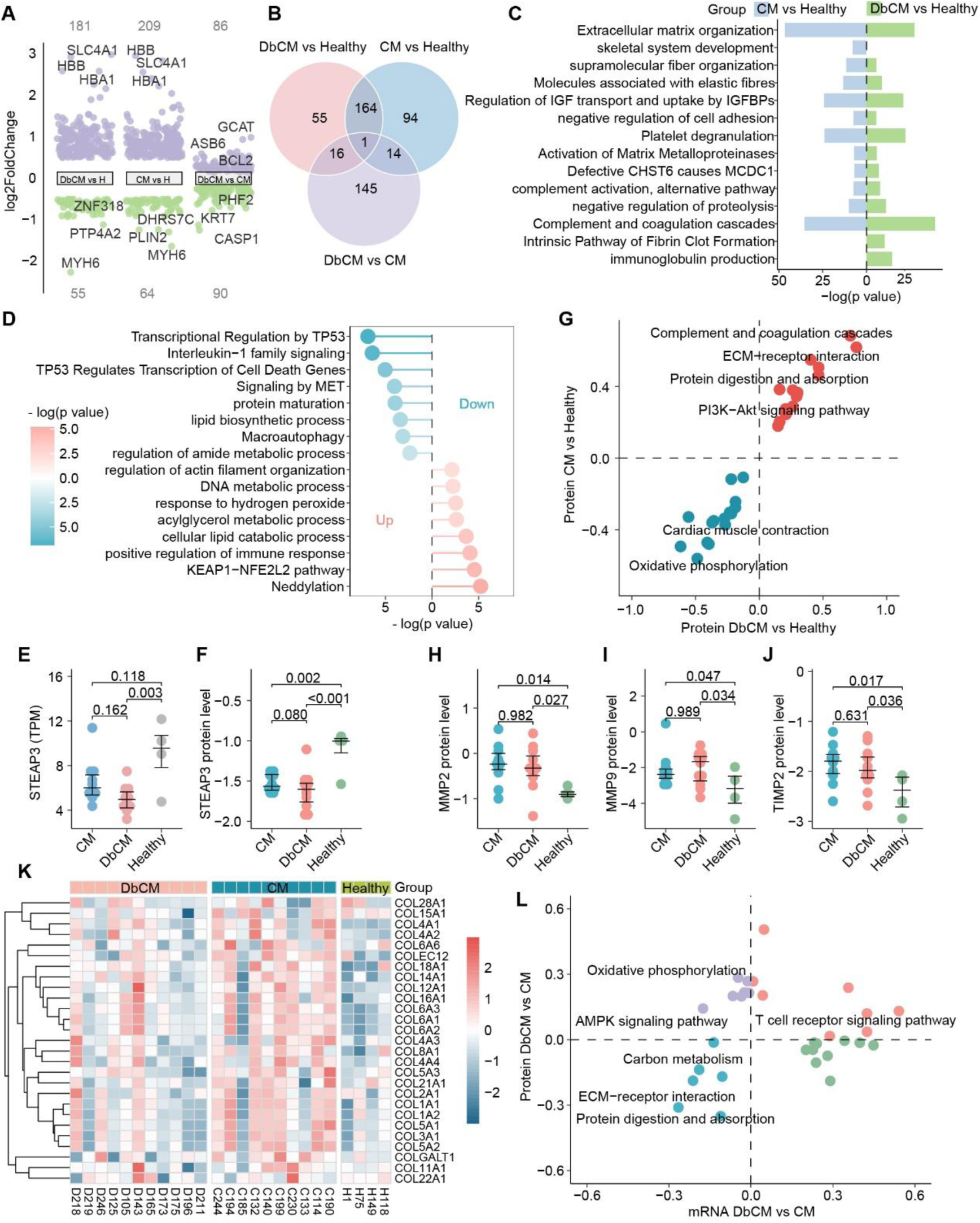
Cardiac fibrosis remodeling was activated in patients. (**A**) Multi-volcano plot of DEPs across multiple comparation groups. (**B**) Venn diagram illustrating the overlap of DEPs. (**C**) Enrichment pathways for DEPs between disease groups and the healthy group, highlighting significant enrichment in pathways related to myocardial fibrosis in both disease groups. (**D**) Metascape enrichment analysis of upregulated and downregulated proteins between DbCM and cardiomyopathy without diabetes groups revealed significant differences in lipid metabolism. (**E, F**) STEAP3 Metalloreductase was specifically downregulated in DbCM patients compared to healthy controls, with significant decreases in both mRNA and protein levels. The horizontal line represents the median and error range. *P*-values from differential analysis were displayed. (**G**) Two-dimensional annotation based on KEGG showing consistent protein alterations in disease groups compared to healthy controls. Each dot represents a KEGG pathway, with the X and Y-axes representing pathway enrichment scores. ECM-receptor interaction was consistently upregulated in both disease groups. (**H-J**) Normalized protein intensities of MMP2, MMP9, and TIMP2. (**K**) Heatmap visualizing collagen protein levels, with collagen levels in DbCM patients showing a less pronounced increase compared to cardiomyopathy patients without diabetes. (**L**) Two-dimensional annotation analysis of transcription and protein alterations between DbCM and cardiomyopathy without diabetes revealed a reduction in ECM-receptor interaction in DbCM patients.

### Cardiac fibrosis remodeling

Myocardial fibrosis is a hallmark of cardiomyopathy [5]. We identified activation of pathways associated with fibrosis initiation, including Wnt and TGF-β signaling (Fig. 2F), along with upregulation of pathways linked to ECM deposition and MMP activation in the disease groups compared to healthy controls (Fig. 3C). Additionally, the ECM-receptor interaction was consistently upregulated in the protein alterations observed across the disease groups relative to the healthy group (Fig. 3G, Source Data 6). These findings suggest that fibrotic remodeling actively occurs in these patients. MMPs, the primary proteases responsible for degrading ECM proteins, along with tissue inhibitors of MMPs (TIMPs), are essential to maintain ECM homeostasis and regulating cardiac remodeling [17]. Gelatinases MMP2 (DbCM: log_2_FC=0.58, *P*=0.027; cardiomyopathy without diabetes: log_2_FC=0.56, *P*=0.014), MMP9 (DbCM: log_2_FC=1.24, *P*=0.034; cardiomyopathy without diabetes: log_2_FC=1.17, *P*=0.047), and inhibitors TIMP2 (DbCM: log_2_FC=0.51, *P*=0.036; cardiomyopathy without diabetes: log_2_FC=0.59, *P*=0.017) were significantly elevated in both DbCM and cardiomyopathy without diabetes groups compared to healthy controls (Fig. 3H-J).

Diabetes in combination with coronary artery disease enhances collagen deposition and exacerbates cardiac fibrosis [18]. However, the increase in collagen levels in patients with DbCM was less pronounced than that in patients with cardiomyopathy without diabetes (Fig. 3K). Moreover, two-dimensional annotation analysis of the transcriptomic and proteomic differences between the two disease groups revealed a reduction in ECM-receptor interactions in patients with DbCM (Fig. 3L). The protein–protein interaction network (Fig. 4A, Source data 7) of DEPs between DbCM and cardiomyopathy without diabetes groups revealed strong interactions among Collagen Type V Alpha 1 Chain (COL5A1, log_2_FC=-0.62, *P*=0.043), Collagen Type V Alpha 2 Chain (COL5A2, log2FC=-0.69, *P*=0.042), and Fibrillin 1 (FBN1, log_2_FC=-0.32, *P*=0.030), all of which were downregulated in DbCM (Fig. 4B-D). Deficiency in Collagen V has been linked to increased scar size following acute myocardial injury, whereas a decrease in FBN1, a pathogenic gene associated with Marfan syndrome, may impair elastic fiber formation and reduce tissue elasticity [19, 20]. To validate the fibrosis proteomic alterations, we performed western blotting and observed the significantly decreased protein levels of MMP2, MMP9, and COL5A1 in DbCM patients compared to cardiomyopathy patients without diabetes (Fig. 4E). Additionally, quantifications of fibrosis showed that the relatively fibrotic areas of DbCM heart tissues were significantly less than cardiomyopathy without diabetes (Fig. 4F-G). In summary, diabetes impairs the matrix repair capacity in patients with cardiomyopathy, potentially leading to decreased cardiac compliance.

**Fig. 4.**
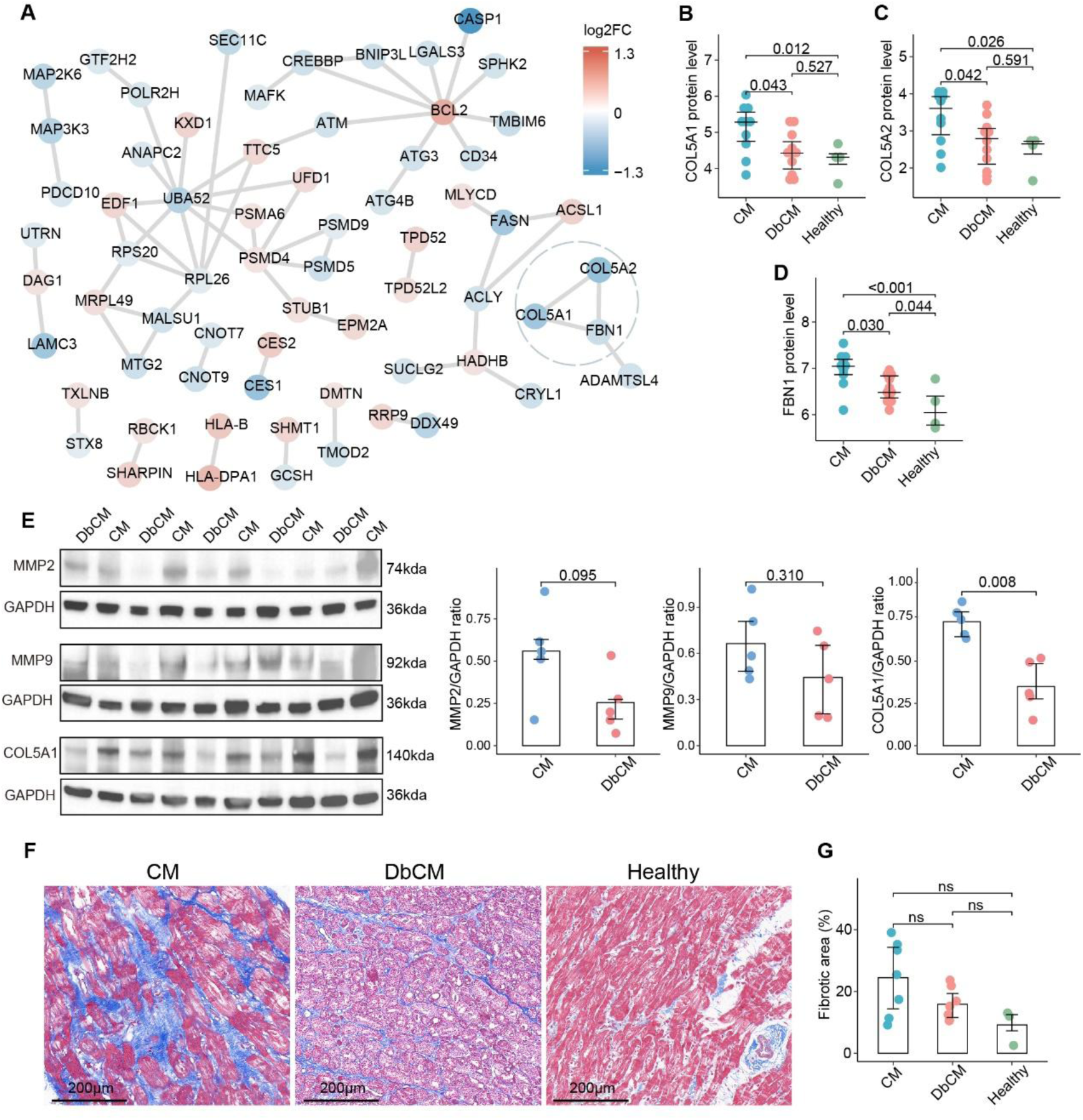
The matrix repair ability of those with diabetes appeared to be relatively impaired. (**A**) Protein-protein interaction (PPI) network of DEPs between DbCM and cardiomyopathy patients without diabetes, constructed using the STRING database with a minimum required interaction score of 0.5, visualized by Cytoscape (v3.10.1). The color of each protein node corresponds log_2_FoldChange value. The PPI network highlights interactions among COL5A1, COL5A2, and FBN1, all of which were downregulated in DbCM compared to cardiomyopathy without diabetes. (**B-D**) Normalized protein intensities of COL5A1, COL5A2, and FBN1, respectively. *P*-values for the differential analysis were displayed. The horizontal line represents the median and error range. (**E**) Protein quantification of human heart tissue samples (DbCM: n=5, cardiomyopathy without diabetes: n=5) for COL5A1, MMP2 and MMP9 by western blotting. (**F**) Masson’s trichrome staining of myocardium from DbCM patients (n=7), cardiomyopathy patients without diabetes (n=7) and healthy donors (n=3). (**G**) Quantification of fibrotic areas. Bars represent mean. Each dot is an average of measurements from 3 fields of view per tissue. Fibrotic differences between groups were accessed by the Mann–Whitney U test, ns: no significant.

### Metabolic alteration landscape

Metabolite levels were significantly altered, with 381 and 536 differentially expressed metabolites (DEMs) identified in the DbCM and cardiomyopathy without diabetes groups, respectively, compared to healthy controls (Fig. 5A-B, Source Data 2). The primary classes of DEMs in both comparisons were glycerophospholipids, amino acids, and their metabolites (Fig. 5C; Source Data 8). Two-dimensional annotation analysis revealed consistent upregulation of protein digestion and absorption pathways in both disease groups compared to the control group in protein alterations (Fig. 3G), with the observed metabolic changes aligned with this finding. In the comparison between the DbCM and cardiomyopathy without diabetes groups, 571 DEMs were identified after adjusting for age, sex, and type of cardiomyopathy (Fig. 5A, Source Data 2). Most downregulated metabolites were triglycerides (TGs), whereas a considerable proportion of upregulated DEMs belonged to glycerophospholipids and fatty acyls (FAs) (Fig. 5C, Source Data 8).

**Fig. 5.**
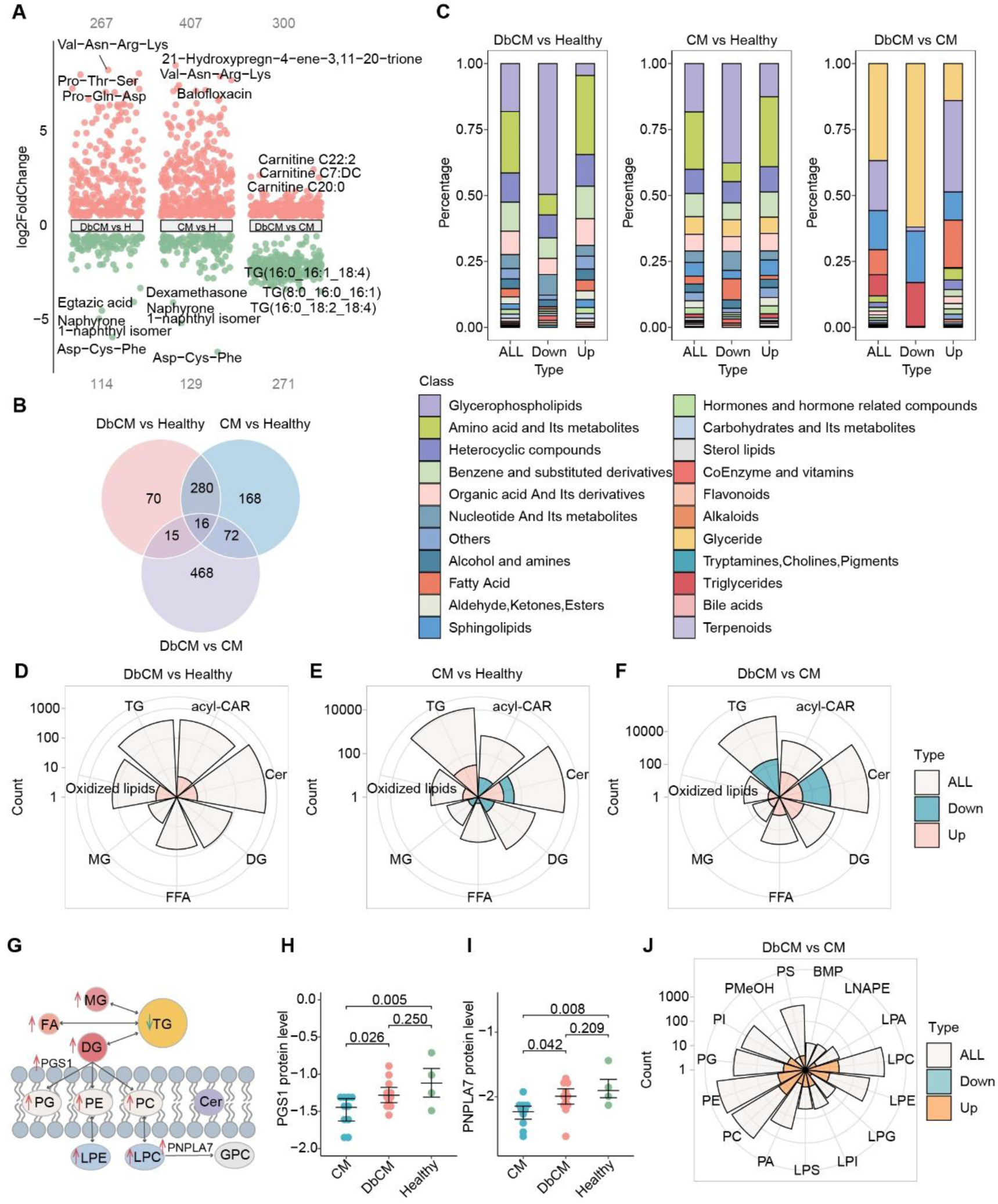
Disruption of myocardial lipid metabolism in DbCM. (**A**) Multi-volcano plot illustrating DEMs. (**B**) Venn diagram showing the overlap of DEMs between the groups. (**C**) Bar chart depicting the distribution of DEMs across categories. Notable alterations were observed in key categories such as glycerophospholipids and amino acids, with lipids being most significantly affected between DbCM and cardiomyopathy without diabetes. (**D-F**) Rose charts representing lipid composition changes. Beige indicates the total number of detected lipids, green shows the number of downregulated lipids, and pink represents upregulated lipids. (**G**) Schematic diagram summarizing significant alterations of glycerophospholipid metabolism identified by the joint enrichment analysis. (**H, I**) Normalized protein intensities of PNPLA7 and PGS1, along with corresponding *P*-values from differential analysis. The horizontal line represents the median and error range. (**J**) Rose chart showing differences in glycerophospholipids between DbCM and cardiomyopathy without diabetes groups.

Lipid accumulation patterns in patients with DbCM differed from those in patients with cardiomyopathy without diabetes and healthy controls (Supplementary Fig. 2). In patients with DbCM, acylcarnitines (acyl-CARs), which are intermediates of fatty acid oxidation (FAO), were upregulated, with no significant changes observed in TG levels (Fig. 5D, Source Data 9). In contrast, patients with cardiomyopathy without diabetes exhibited elevated TGs and oxidized lipid levels, whereas acyl-CARs were reduced (Fig. 5E). A comparison between patients with DbCM and cardiomyopathy without diabetes revealed that all TGs were significantly downregulated in the DbCM group, whereas acyl-CARs, monoglycerides, and diglycerides were upregulated (Fig. 5F). Furthermore, glycerophospholipid metabolism was significantly enriched in the analysis of DEPs and DEMs between disease groups (Fig. 5G, Supplementary Table 2). Elevated levels of lysophospholipase patatin-like phospholipase domain-containing 7 (PNPLA7) and phosphatidylglycerophosphate synthase (PGS1) were also observed (Fig. 5H-I). In the DbCM group, the DEMs of phosphatidylcholines, lysophosphatidylcholines, phosphatidylethanolamines, lysophosphatidylethanolamines, and phosphatidylglycerols were upregulated compared to those in the cardiomyopathy without diabetes group (Fig. 5J).

### Increased utilization of triglyceride-derived fatty acids by the heart in DbCM

A healthy adult heart exhibits high metabolic adaptability and flexibility, allowing for efficient switching between different energy substrates to maintain energy supply [21]. Metabolomic analysis revealed that FAO was reduced in cardiomyopathy patients without diabetes and increased in patients with DbCM, as indicated by contrasting alterations in TGs, the source of fatty acids, and the oxidation intermediate acyl-CARs between the two groups (Fig. 5D-E). However, no significant differences were observed in the levels of proteins involved in fatty acid transport, such as CD36 and fatty acid-binding proteins, between the patient groups. DEPs between the DbCM and cardiomyopathy without diabetes groups were upregulated in acylglycerol metabolism, but downregulated in lipid biosynthesis (Fig. 3D). Fatty acid synthase (FASN), a key enzyme in the de novo synthesis of fatty acids, was significantly downregulated in patients with DbCM compared to cardiomyopathy patients without diabetes (log_2_FC=-0.60, *P*=0.047) (Fig. 6A), suggesting that the primary source of fatty acids for myocardial FAO in DbCM is TG breakdown. Furthermore, acyl-CoA synthetase long-chain family member 1 (ACSL1) levels were elevated in the DbCM group (log_2_FC=0.32, *P*=0.010) (Fig. 6B), indicating increased oxidative utilization of fatty acids. In summary, elevated levels of fatty acids and acyl-CARs, coupled with reduced TG levels in patients with DbCM compared to those in cardiomyopathy patients without diabetes, indicate enhanced FAO in DbCM (Fig. 6C). The widespread decrease in TG levels suggests an increased utilizations on TG in hearts of DbCM.

**Fig. 6.**
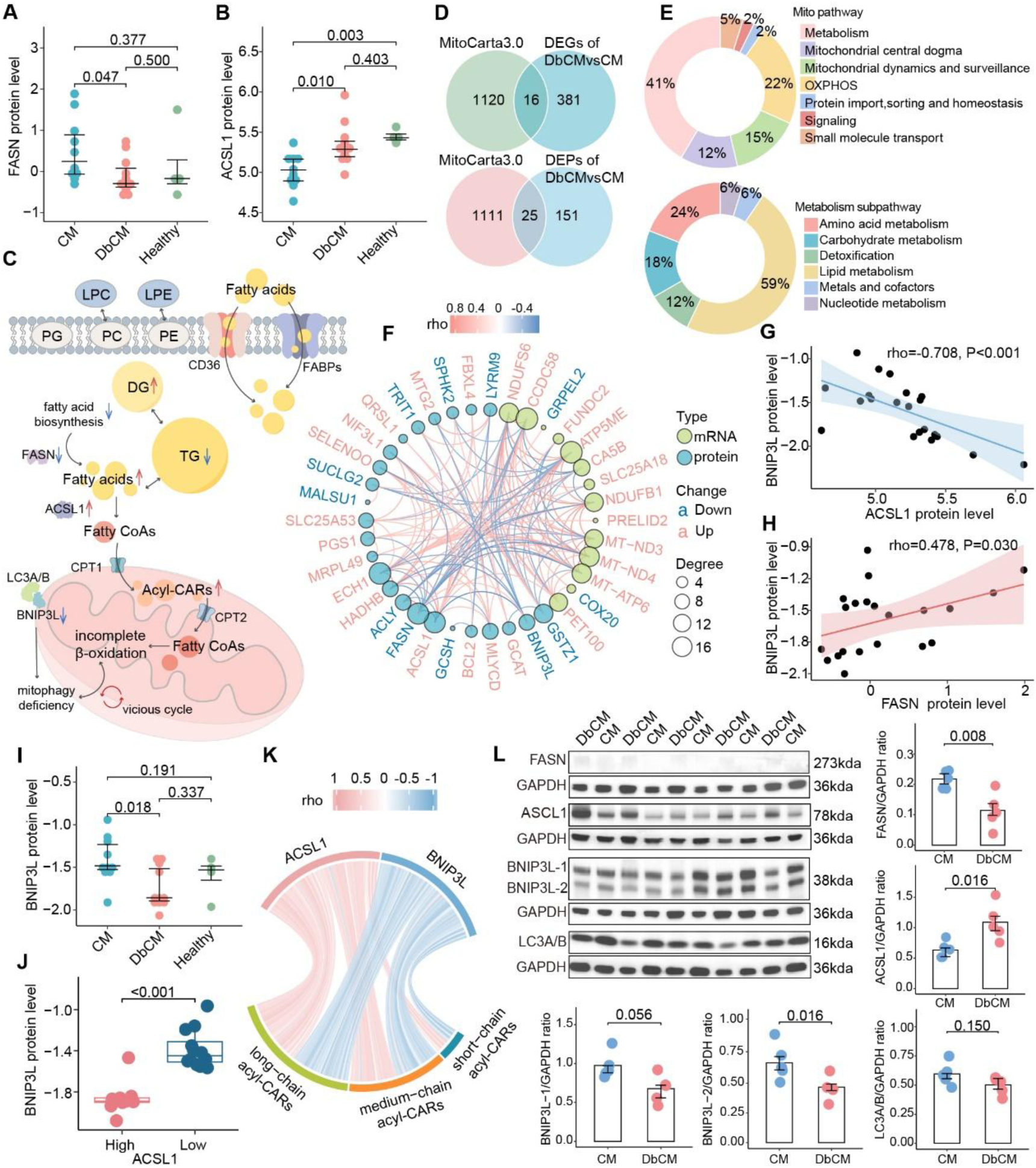
Mitophagy in diabetes hearts was impaired and closely related to fatty acid oxidation. (**A, B**) Normalized proteins intensities of FASN and ACSL1 were presented. *P*-values derived from differential analysis are indicated. The horizontal line represents the median, with the error range shown. (**C**) A schematic illustrating the sources and utilization of fatty acids. No significant changes were detected in fatty acid transporters, while biosynthesis was suppressed, and incomplete β-oxidation was upregulated, leading to the accumulation of acyl-CARs. This incomplete β-oxidation, combined with defective mitophagy, initiates a detrimental feedback loop. (**D**) Venn diagrams showing the overlap of 41 genes and proteins between DEGs and DEPs and the MitoCarta3.0 dataset. The intersecting sets were categorized as MitoDEGs and MitoDEPs. (**E**) Mitochondrial pathways associated with MitoDEGs and MitoDEPs, with percentages representing the proportion of genes and proteins involved in each pathway. Sub-pathways of mitochondrial metabolism were further detailed. (**F**) Correlation network of MitoDEGs and MitoDEPs, with Spearman correlation coefficients indicated. Significantly upregulated and downregulated genes and proteins were colored in pink and blue, respectively. The degree of each node represents its number of connections. (**G, H**) Scatter plots depicting the correlations between ACSL1 and BNIP3L protein levels, as well as FASN and BNIP3L. (**I**) Normalized protein intensities of BNIP3L, showing a significant reduction in DbCM compared to cardiomyopathy without diabetes. *P*-values from differential analysis were provided. (**J**) Patients were grouped based on the median protein level of ACSL1, with the high-ACSL1 group exhibiting significantly lower BNIP3L levels. (**K**) A chord diagram illustrating the correlations between the protein levels of BNIP3L and ACSL1 and the levels of acyl-CARs. (**L**) Protein quantification of human heart tissue samples (DbCM: n=5, cardiomyopathy without diabetes: n=5) for FASN, ACSL1, BNIP3L and LC3A/B by western blotting. Bars represent mean. Two-sided *P*-values were estimated using the Mann-Whitney U test.

### Mitophagy was impaired and related to FAO

Mitochondrial quality control is a critical area of investigation in DbCM [22]. We further examined the mitochondrial functions of DEGs and DEPs between patients with DbCM and cardiomyopathy without diabetes. We identified 16 MitoDEGs and 25 MitoDEPs that overlapped in the dataset (Fig. 6D, Source Data 10). Of these, 41% (n=17) were linked to metabolism (Fig. 6E), primarily lipid and amino acid metabolism. Additionally, 22% (n=8) were involved in oxidative phosphorylation, and 15% (n=6) were associated with mitochondrial dynamics and surveillance, including mitophagy (Fig. 6E). In the correlation network of MitoDEGs and MitoDEPs (Fig. 6F), the levels of BCL2 interacting protein 3-like (BNIP3L, also known as NIX) in patients showed a significant negative correlation with ACSL1 (rho=-0.708, *P*<0.001) and a positive correlation with FASN (rho=0.478, *P*=0.030) (Fig. 6G-H). BNIP3L, a key mitophagy inducer, was significantly reduced in patients with DbCM compared to those with cardiomyopathy without diabetes (log_2_FC=-0.30, *P*=0.018) (Fig. 6I).

Mitophagy dysregulation is a key mechanism in various metabolic and cardiovascular diseases [22]. Furthermore, when patients were categorized based on median ACSL1 protein expression, those in the high ACSL1 expression group exhibited significantly lower levels of BNIP3L (Fig. 6J). Metabolic correlation analysis revealed a positive correlation between ACSL1 protein levels and acyl-CARs, whereas BNIP3L levels were negatively correlated with acyl-CARs (Fig. 6K, Source Data 11). Protein quantifications validated that levels of ACSL1 were significantly increased in DbCM hearts compared to cardiomyopathy without diabetes (Fig. 6L). Additionally, protein levels of BNIP3L and microtubule-associated protein 1-light chain 3 (LC3A/B), which have central role in autophagosome formation, were significantly decreased in DbCM (Fig. 6L). Our results indicated that patients with DbCM exhibited mitophagy dysfunction mediated by BNIP3L, which was significantly correlated with multiple aspects of fatty acid metabolic dysregulation, in contrast to patients with cardiomyopathy without diabetes.

### Multi-omics integration

Unbiased deconstruction of extensive variability across multilayer omics data and identification of diabetes-relevant alterations are essential for gaining deeper insights. Bidirectional orthogonal partial least squares models revealed cross-omics correlations, suggesting that the dysregulation of oxidative phosphorylation, fatty acid metabolic processes, and the cytoskeleton of muscle cells was significantly influenced by multiple omics factors (Supplementary Fig. 3A-E, Source Data 12). To explore clinical relevance, we integrated three omics datasets from 21 patients using multi-omics factor analysis. This approach allowed us to derive a model comprising five factors that collectively explained 48.2%, 13.0%, and 65.7% of the variance in the transcriptome, proteome, and metabolome, respectively (Fig. 7A, Source Data 13). Factors 2 (rho=0.457, *P*=0.037) and 3 (rho=0.535, *P*=0.012) exhibited significant positive correlated with diabetes (Fig. 7D). Lipids-related features showed a strong association with diabetes, with ACSL1 being identified as the top-weighted protein in Factor 3 (Fig. 7C). Enrichment analysis of the top 20 proteins and genes, based on the absolute values of their weights, revealed that immune- and ECM-related pathways and lipid metabolism were strongly associated with DbCM (Fig. 7D).

**Fig. 7.**
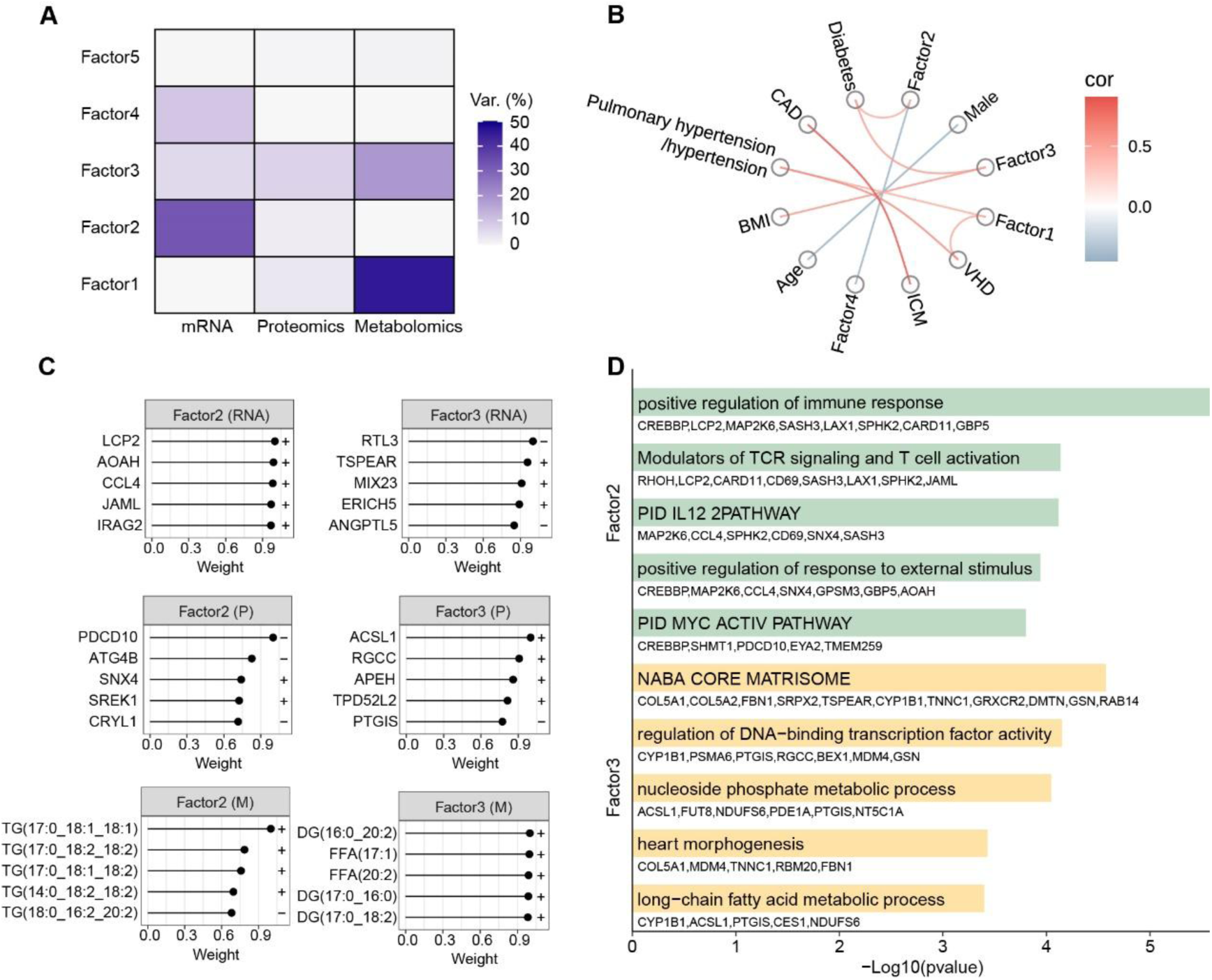
Multi-omics integration revealed significant disturbances in lipid metabolism. (**A**) Multi-omics data were subjected to dimensionality reduction, resulting in a model comprising five factors, which collectively explained 48.2% of the transcriptome variance, 13.0% of the proteome variance, and 65.7% of the metabolome variance. (**B**) Spearman correlation network illustrating the correlations between factors and clinical characteristics, with factor 2 and factor 3 showing a significant positive correlation with diabetes. (**C**) Top five weighted genes, proteins and metabolites of factor 2 and factor 3. Symbols indicate the direction (positive or negative) of the weights. (**D**) Enriched pathways for the top 20 proteins and genes, based on the absolute values of their weights, highlight the significant roles of immune-related pathways, cellular matrix pathways, and lipid metabolism in DbCM.

### Mouse mutation models for key proteins

To compensate for the lack of direct *in vivo* validation and to clarify the potential physiological roles of the key proteins ACSL1, FASN, BNIP3L, COL5A1, COL5A2, FBN1, and STEAP3, we analyzed genetically modified mouse models using data from the Mouse Genome Informatics database. The objective was to determine whether alterations in these genes are linked to phenotypes associated with DbCM. Mutations associated with cardiovascular abnormalities were also identified. Transgenic models with inserted expressed sequences for ACSL1 exhibited phenotypes, including cardiomyopathy, increased cardiac muscle triglyceride levels, abnormal myocardial fiber morphology, and reduced heart ventricle muscle contractility. Knockout models for other genes displayed phenotypes such as impaired mitophagy, altered left ventricular morphology, cardiac fibrosis, disrupted collagen fibril structure, and impaired glucose tolerance (Supplementary Table 3). Importantly, these proteins also served as effective biomarkers in distinguishing DbCM from cardiomyopathy without diabetes in human patients (Supplementary Fig. 4). These findings highlight the intrinsic roles of these genes in the regulation of heart function.

## Discussion

Our analysis provided several key insights. First, we identified distinct perturbations in lipid metabolism and mitochondrial function in patients with DbCM compared to those in patients with cardiomyopathy without diabetes. Second, hearts of individuals with DbCM exhibited increased utilization of triglyceride-derived fatty acids for energy production. Furthermore, BNIP3L-mediated mitophagy impairment was more pronounced in DbCM. The correlation between BNIP3Lprotein levels and those of ACSL1 and FASN suggests that compromised mitochondrial quality control is associated with dysregulated fatty acid metabolism. Finally, although myocardial fibrosis was evident in both groups, DbCM hearts demonstrated a relatively impaired matrix repair capacity for extracellular matrix remodeling, specifically evidenced by reduced COL5A1 protein levels. Collectively, these findings provide novel mechanistic insights into diabetes-associated cardiac dysfunction and may inform future therapeutic development.

In cardiac conditions such as cardiomyopathy or HF, the myocardium reduces or maintains its reliance on FAO owing to the inefficiency of the energy supply [23]. However, in patients with obesity or diabetes, FAO often increases, which is driven by a high-glucose environment and insulin resistance [24]. Our study identified distinct metabolic patterns between patients with DbCM and those without diabetes. Specifically, we observed an increase in FAO in patients with DbCM, and a decrease in the number of patients with cardiomyopathy without diabetes. By comparing these groups, we found that the de novo fatty acid synthesis pathway in DbCM hearts was downregulated, whereas triglyceride reserves were significantly reduced. FAs, especially acyl-CARs, and acyl-CoA synthesis were notably upregulated in DbCM hearts. No significant differences in free fatty acid transporters were observed, suggesting that the free fatty acids used by the DbCM heart were primarily derived from TG breakdown, accompanied by the production of monoglycerides and diglycerides, which are linked to lipotoxicity and insulin signaling interference [5, 23]. The critical differences between DbCM and cardiomyopathy patients without diabetes are the marked increase in fatty acid utilization and incomplete oxidation in DbCM hearts. We observed a significant increase in ACSL1 protein levels in patients with DbCM, which correlates with changes in fatty acid utilization and human heart tissue [13, 25, 26]. ACSL1 has also been implicated in loss of myocardial regenerative capacity [27]. Furthermore, acyl-CARs, which are the intermediates of incomplete fatty acid oxidation, serve as markers of mitochondrial FAO dysfunction in DbCM [13, 25, 28]. Elevated circulating acyl-CAR levels have also been reported in patients with DbCM [29, 30]. In summary, the DbCM heart demonstrated a metabolic shift, characterized by an increased reliance on TG breakdown for fatty acid supply, along with heightened FAO to meet energy demands. However, this shift results in the accumulation of lipotoxic intermediates and incomplete FAO, contributing to cellular dysfunction.

Mitochondrial dysfunction is a key characteristic of DbCM, in which damaged mitochondria undergo mitophagy, a process that degrades structurally compromised mitochondria via the PINK1-PRKN/PARKIN pathway and receptor-mediated pathways involving proteins such as BNIP3, BNIP3L, and FUNDC1 [9, 31, 32]. Impaired mitophagy leads to the accumulation of damaged mitochondria and lipids, exacerbating DbCM [33]. We observed a significant reduction in BNIP3L protein levels in patients with DbCM compared with those in individuals with cardiomyopathy without diabetes. Similar downregulation of BNIP3L has been shown in myocardial samples from patients with HF due to ischemic or dilated cardiomyopathy compared to healthy controls, and inhibition of BNIP3L-mediated mitophagy in adult cardiac progenitor cells leads to mitochondrial fission and the formation of damaged mitochondria [34, 35]. Furthermore, the activation of mitophagy protects against DbCM induced by a high-fat diet, whereas BNIP3L activation can protect retinal pigment epithelial cells from high glucose-induced injury [36, 37]. These findings suggest that diabetes exacerbates mitophagy defects by inhibiting the BNIP3L pathway, thus worsening mitochondrial dysfunction and contributing to DbCM progression. Additionally, we found significant correlations between BNIP3L and fatty acid metabolism-related proteins, including ACSL1, FASN, and acyl-CARs, indicating a close link between fatty acid metabolism and receptor-mediated mitophagy. Increased FAO and impaired mitophagy are key features of DbCM [1, 5]. Although it was challenging to confirm a causal relationship between these processes, a mutual exacerbation was evident. The deletion of BNIP3L leads to the accumulation of hypoxia-inducible factors, which decrease FASN expression and disrupt long-chain FAO, resulting in metabolic defects and impaired adenosine-triphosphate synthesis [38]. In summary, BNIP3L-mediated mitophagy is crucial for maintaining mitochondrial homeostasis and metabolic regulation in hearts with DbCM. Its influence on fatty acid metabolism and oxidative balance positions BNIP3L as a potential therapeutic target for the treatment of DbCM.

ECM remodeling plays a pivotal role in the pathogenesis of DbCM; however, the impact of diabetes on cardiac matrix repair remains underexplored [1, 5, 18, 39]. ECM is essential to provide structural support and regulate critical cellular functions [40]. We observed a significant downregulation of COL5A1, COL5A2, and FBN1 proteins in patients with DbCM compared to that in cardiomyopathy patients without diabetes, along with a reduction in ECM-receptor interactions. Collagen V deficiency is linked to an increased scar size and disrupted wound healing after acute heart injury [19]. In COL5A1 knockout mice, reduced scar stiffness and compromised cardiac contractile forces have been observed, with myofibroblast differentiation driven by altered mechanosensitive receptor integrin expression in cardiac fibroblasts [19]. Additionally, FBN1, a microfibrillar protein crucial for structural integrity and elastic strength, was reduced in DbCM, impairing elastic fiber formation and contributing to tissue laxity [20]. FBN1 expression is influenced by H3 epigenetic modifications in hyperglycemic/insulin-resistant rats, with reduced mRNA levels in the myocardial tissue [41]. Notably, elevated FBN1 expression promotes fibroblast proliferation and migration, while inhibiting apoptosis and facilitating wound healing in diabetic foot ulcers [42]. Additionally, we discovered that STEAP3, a novel potential target in DbCM, was specifically downregulated at both the mRNA and protein levels in patients with DbCM. This decrease in STEAP3 expression may impair ECM deposition and remodeling by affecting iron levels in diabetic murine wounds, thus hindering wound recovery [43, 44]. In summary, the cardiac remodeling mechanisms in DbCM differ significantly from those in patients with diabetes. Diabetes may impair the matrix repair capacity of DbCMs, thereby reducing cardiac and vascular compliance. Therefore, antifibrotic therapies should be approached with caution, ensuring that the matrix repair ability is preserved, as premature interventions may lead to adverse outcomes [39]. Insights from diabetic foot ulcer wound healing and keratin repair research may provide valuable strategies for the prevention and repair of matrix damage in DbCMs.

In our study, patients with DbCM and those with cardiomyopathy without diabetes exhibited downregulated immune and inflammatory regulation compared to healthy controls, a finding consistent with the validation analysis. This observation may help explain the largely unsuccessful outcomes of clinical trials testing anti-inflammatory therapies for HF, highlighting the significant heterogeneity in the inflammatory profiles of HF patients [45, 46, 47]. Additionally, this downregulation may reflect immune exhaustion in end-stage HF or the effects of preoperatively administered immunosuppressive therapies. These findings underscore the complexity of inflammatory regulation in HF and suggest that alterations in the immune system contribute to disease progression.

Although this study offers the most comprehensive multi-omics characterization of DbCM in the human heart, it has some limitations. First, the relatively small sample size dictated by restricted tissue accessibility may have affected the statistical robustness of our findings. Second, because the patient samples were derived from end-stage transplanted hearts, the observed molecular alterations may not be exclusive to myocardial cells and may represent late-stage pathophysiological changes, complicating the establishment of causal relationships. Third, although this study highlights several novel insights, these findings have not yet been validated through *in vitro* or *in vivo* experiments and should be interpreted with caution. Due to the limited availability of cardiac data, the external validation datasets referenced in the transcriptome comparison do not fully align with the sample characteristics of this cohort, which may restrict the interpretability and generalizability of the validation results. Additionally, while the clinical impact of emerging glucose-lowering therapies such as SGLT2 inhibitors is an important direction for future research, limited drug exposure within the cohort precludes relevant analyses. Addressing these limitations in future investigations will be crucial to refine the mechanistic understanding and therapeutic strategies for DbCM.

In conclusion, this study presents a comprehensive multi-omics characterization of human left ventricular tissues in DbCM, revealing pathophysiological distinctions from cardiomyopathy without diabetes. We identified profound alterations in lipid metabolism, mitochondrial function, and extracellular matrix remodeling in the hearts of patients with DbCM. Our findings also highlighted a potential mechanistic link between BNIP3L-mediated mitophagy deficiency and dysregulated fatty acid metabolism during DbCM progression. These insights advance our understanding of the molecular insights of diabetes-associated cardiac dysfunction and may inform the development of targeted therapeutic strategies.

## Methods

### Human heart tissue collection

Left ventricular tissues were collected from patients diagnosed with ischemic or dilated cardiomyopathy who underwent cardiac transplantation at Guangdong Provincial People’s Hospital, Guangzhou, China, between August 2019 and April 2022. Patients with a confirmed diagnosis of T2D were classified as having diabetic cardiomyopathy (DbCM group, n=11), whereas patients without diabetes were assigned to the cardiomyopathy without diabetes group (CM group, n=11). The two groups were matched for age (±4 years) and sex in a case-control study. In addition, healthy myocardial tissues (healthy group, n=4) were obtained from donor hearts that could not be transplanted in time. All excised heart tissues were immediately flash-frozen and stored in liquid nitrogen until further analysis. This study was conducted in accordance with the Declaration of Helsinki and approved by the Ethics Committee of Guangdong Provincial People’s Hospital (GDREC2016255H). Written informed consent was obtained from all participants.

### RNA-sequencing

Total RNA was extracted from frozen tissues using the RNeasy Plus Micro Kit (Qiagen) following the manufacturer’s protocol. RNA quality and integrity were assessed using an Agilent 2100 Bioanalyzer (Agilent Technologies). RNA sequencing libraries were prepared with the NEBNext Ultra Directional RNA Library Prep Kit for Illumina (NEB, E7420) and sequenced on an Illumina NovaSeq 6000 platform. Clean reads were generated by removing adapter sequences, poly-N stretches, and low-quality reads. High-quality paired-end reads were aligned to the human reference genome (GRCh38, release 109) using Hisat2 (v2.0.5), achieving mapping rates exceeding 96%. Gene expression was quantified using StringTie (v1.3.3b), and genes absent in more than one-third of the samples were excluded. Detailed processing steps are provided in the Supplementary Information.

### 4D-DIA proteomics analysis

Quantitative proteomic analysis was performed using 4D data-independent acquisition (4D-DIA). Raw mass spectrometry data were processed with DIA-NN (v1.8.1) in a library-free mode. A spectral library was generated using the Homo sapiens SwissProt database (20,376 entries) via deep neural network prediction, and refined using match-between-runs (MBR) strategies. The false discovery rate (FDR) was controlled at <1% at both the protein and precursor ion levels. Protein intensities were median-normalized across samples. Proteins missing in more than one-third of the samples were excluded, and missing values were imputed using the k-nearest neighbors (KNN) method. Full processing details are provided in the Supplementary Information.

### Metabolite extraction and full-spectrum widely targeted metabolomics data processing

Both hydrophilic and hydrophobic metabolites were extracted to maximize the coverage. The raw data were processed using MultiQuant (v3.0) for chromatographic peak integration and calibration. Metabolite intensities were quantified based on peak areas. Metabolites missing in more than one-third of the samples were excluded and missing values were imputed using the KNN method. Quality control was performed using pooled quality control samples spiked with internal standards. Metabolite identification was based on public and in-house databases, including the Human Metabolite Database and KEGG. Further details are provided in the Supplementary information.

### Clinical data analysis

Statistical analyses were performed using R software (v4.2.2). Clinical characteristics were compared using the Mann–Whitney U test for continuous variables and Fisher’s exact test for categorical variables. A two-sided significance threshold of *P*<0.05 was applied.

### Omics data processing and differential analysis

The transcriptomic read counts were normalized to the number of transcripts per million. The metabolomic and proteomic intensity matrices were log_2_-transformed prior to analysis. Principal component analysis and PLS-DA were used to visualize group separation across the datasets.

Differential expression thresholds for statistical significance and effect size were adapted based on biological priors, platform-specific sensitivities, and inherent molecular properties [28, 48]. DEGs were identified using DESeq2 with FDR adjustments using the Benjamini–Hochberg method. DEGs were defined as follows: 1) DbCM vs. healthy and cardiomyopathy without diabetes vs. healthy, FDR<0.1 and |log_2_ fold change|>0.5; 2) DbCM vs. cardiomyopathy without diabetes, |log_2_ fold change|>0.5 and *P*<0.05, adjusted for age, sex, and cardiomyopathy type. Limma was used to identify DEPs and DEMs across the groups. For DbCM vs. healthy or cardiomyopathy without diabetes vs. healthy, proteins and metabolites with *P*<0.05 and |log_2_ fold change|>0.5 were classified as upregulated or downregulated DEPs/DEMs, respectively. For DbCM vs. cardiomyopathy without diabetes, proteins with *P*<0.05, adjusted for age, sex, and type of cardiomyopathy, and metabolites with |log_2_ fold change|>0.5 and *P*<0.05, adjusted for age, sex, and type of cardiomyopathy, were considered DEPs and DEMs for subsequent analysis.

Bidirectional orthogonal partial least squares analysis was used to assess inter-omics associations between DEGs, DEPs, and DEMs [49]. Multi-omics integration was performed using multi-omics factor analysis across omics profiles of samples [50].

### Functional and pathway analysis

To investigate mitochondrial function, MitoCarta3.0 was used to identify mitochondrial-related DEGs (MitoDEGs) and DEPs (MitoDEPs) [51]. GSEA was performed on the transcriptome using KEGG pathway annotations and ClusterProfiler package. Proteomic functional analysis was conducted using Metascape [52]. Joint pathway enrichment of metabolomics with proteomic data was analyzed using MetaboAnalyst [53]. Two-dimensional annotation analysis using Perseus was employed to compare the consistency of KEGG pathway changes in the transcriptome and proteome [54]. Immune and stromal cell infiltration was quantified using CIBERSORT and xCell [55, 56]. Protein-protein interaction networks were constructed based on STRING database predictions. The Mouse Genome Informatics database was used to investigate the phenotypic associations of candidate genes with cardiovascular or metabolic traits [57]. The discriminatory performance of key molecules distinguishing DbCM from cardiomyopathy without diabetes was evaluated using the area under the curve.

### External validation of transcriptomic findings

External validation was performed by analyzing the publicly available Gene Expression Omnibus dataset GSE26887, which included transcriptomic profiles of the left ventricular myocardium from 19 HF patients (seven with diabetes, 12 without) and five non-HF controls [15]. Functional and immune cell infiltration analyses were repeated for cross-validation.

### Histological assessments

Left ventricular tissue samples were fixed in 4% paraformaldehyde for 24 h at room temperature. Following fixation, tissues were embedded in paraffin and sectioned at a thickness of 5 μm. After deparaffinization and rehydration, the adjacent sections were stained with Masson’s trichrome to visualize collagen deposition associated with fibrosis. Images were captured using an Eclipse E100 microscope (Nikon, Tokyo, Japan). Fibrotic areas were quantified using Fiji software by calculating the ratio of the blue-stained (collagen) area to the total tissue area.

### Western blotting

Proteins were extracted by centrifuging tissue lysates at 1,000 ×g for 15 minutes at 4°C, and the supernatant was collected. Protein concentrations were determined using the bicinchoninic acid protein assay kit (Thermo Fisher Scientific). Equal amounts of protein (∼30 μg) were separated via 10% sodium dodecyl sulfate-polyacrylamide gel electrophoresis and transferred onto 0.45-μm nitrocellulose membranes (Millipore). The membranes were blocked with 5% bovine serum albumin in Tris-buffered saline containing 0.1% Tween-20 for 2 h at room temperature. After blocking, the membranes were incubated overnight at 4°C with primary antibodies against the indicated proteins, followed by incubation with horseradish peroxidase-conjugated secondary antibodies for 2 h at room temperature. Protein bands were detected using an enhanced chemiluminescence system (GE Healthcare) according to the manufacturer’s instructions. Densitometric quantification of the western blot bands was performed using ImageJ software. The following primary antibodies were used: anti-ACSL1 (4047; Cell Signaling Technology), anti-BNIP3L (12396; Cell Signaling Technology), anti-FASN (3180; Cell Signaling Technology), anti-LC3A/B (12741; Cell Signaling Technology), anti-MMP2 (ab92536; Abcam), anti-MMP9 (ab76003; Abcam), and anti-COL5A1 (Santa Cruz Biotechnology, sc-133162).

## Supporting information

Supplementary Table 1 Clinical information of patients

Supplementary Table 2 metabolomics enrichment results

Supplementary Table 3 MGI annotation

## Acknowledgments

The chief acknowledgment is to the participants in this research and all clinic and research staff for running this study. This study received support from the Natural Science Foundation of China (12331017, 12271118,82203249), Guangdong Basic and Applied Basic Research Foundation (2024A1515013260, 2023A1515010110, 2025A1515012707), the Shenzhen Science and Technology Program (ZDSYS20230626091203007).

## Author contributions

Conceptualization: Q.H.L, S.L.T, J.F, Y.H.W, G.Z.J

Methodology: Q.H.L, S.L.T, L.C, M.T.L, L.G.J, J.S.Y, J.F, Y.H.W, G.Z.J

Data curation: Q.H.L, S.L.T, Y.W.W, L.C, M.T.L, J.Q.C

Investigation: Q.H.L, S.L.T, Y.W.W, L.C, M.T.L, S.J.F, J.Q.C, P.J.W, L.G.J, J.S.Y, J.F, Y.H.W, G.Z.J

Formal analysis: Q.H.L, S.L.T, Y.W.W, M.T.L, P.J.W

Resources: S.L.T, P.J.W, J.F, Y.H.W, G.Z.J

Validation: Q.H.L, S.L.T, Y.W.W, M.T.L, S.J.F, P.J.W, L.G.J

Visualization: Q.H.L, Y.W.W, M.T.L,

Writing—original draft: Q.H.L, S.L.T

Writing—review & editing: Q.H.L, S.L.T, Y.W.W, J.F, Y.H.W, G.Z.J

Supervision: S.L.T, J.F, Y.H.W, G.Z.J

Project administration: Q.H.L, S.L.T, J.F, Y.H.W, G.Z.J

