## Supplementary materials for "Multi-omics reveals lipid metabolism, mitochondrial and extracellular matrix dysregulation of diabetic cardiomyopathy in the human heart"

**Description of Supplementary Information：**

Supplemental methods: Details of multi-omics data processing

Supplementary Table 1: Clinical information of patients

Supplementary Table 2: Metabolomics enrichment results

Supplementary Table 3: Cardiovascular and metabolic homeostasis phenotype information of key proteins from the Mouse Genome Informatics (MGI) database

Supplementary Figure 1: Overview of multi-omics data from human heart tissues.

Supplementary Figure 2 Box plots of global lipid abundance.

Supplementary Figure 3 O2PLS models revealed reveal cross-omics correlations.

Supplementary Figure 4 ROC curves of selected key proteins.

Supplementary Figure 5 Unprocessed, uncropped scans of all blots.

**Supplemental methods**

**RNA Sequencing and Data Processing**

Total RNA was extracted from frozen left ventricular tissues using the RNeasy Plus Micro Kit (Qiagen) according to the manufacturer's instructions. RNA quality and integrity were assessed with the Agilent 2100 Bioanalyzer (Agilent Technologies). RNA sequencing libraries were prepared using the NEBNext Ultra II Directional RNA Library Prep Kit for Illumina (NEB, E7420) following the manufacturer's protocol. Ribosomal RNA (rRNA) was removed from total RNA samples, and fragmentation was performed using divalent cations at elevated temperatures in the First Strand Synthesis Reaction Buffer. First-strand complementary DNA (cDNA) synthesis was carried out using random hexamer primers and M-MuLV Reverse Transcriptase (RNase H-), followed by second-strand cDNA synthesis with DNA Polymerase I and RNase H. Residual overhangs were converted into blunt ends through exonuclease and polymerase activities. After adenylation of the 3’ ends, NEBNext adaptors with hairpin loop structures were ligated to the cDNA fragments. Size selection for 370-420 bp fragments was performed using the AMPure XP system (Beckman Coulter). The adaptor-ligated cDNA was treated with 3 µL USER enzyme (NEB) at 37°C for 15 min, followed by incubation at 95°C for 5 min prior to PCR amplification. Libraries were amplified using Phusion High-Fidelity DNA polymerase, Universal PCR primers, and Index primers, and subsequently purified using the AMPure XP system. Final library quality was assessed using the Agilent 5400 system and quantified by qPCR (final concentration 1.5 nM). Libraries passing quality control were pooled and sequenced on an Illumina NovaSeq 6000 platform with paired-end reads.

Raw sequencing reads were processed to remove adapter contamination, poly-N stretches, and low-quality reads, yielding clean reads for downstream analysis. The reference genome index (GRCh38, release 109) was built using HISAT2 (v2.0.5), and clean paired-end reads were aligned to the reference genome with a mapping rate exceeding 96% across all samples. Gene expression quantification was performed using StringTie (v1.3.3b), and genes missing read counts in more than one-third of the samples were excluded from subsequent analyses.

**Proteomics Sample Preparation and 4D-DIA Analysis**

Myocardial tissue samples were ground in liquid nitrogen and lysed in buffer containing 7 M urea, 2 M thiourea, 4% SDS, and 40 mM Tris-HCl (pH 8.5), supplemented with 1 mM PMSF and 2 mM EDTA. After 5 min incubation, 10 mM DTT was added, and samples were sonicated on ice for 5-15 min. Lysates were centrifuged at 13,000 rpm for 20 min at 4°C, and the supernatants were precipitated with four volumes of pre-cooled acetone at -20°C for 2 h. Pellets were air-dried and resuspended in 8 M urea/100 mM TEAB (pH 8.0). Proteins were reduced with 10 mM DTT at 56°C for 30 min and alkylated with 50 mM iodoacetamide (IAM) at room temperature for 30 min in the dark. Following a second acetone precipitation, protein concentrations were determined using a BCA assay. Equal amounts (~100 μg) of protein from each sample were digested with trypsin at a 1:50 enzyme-to-substrate ratio (w/w) at 37°C overnight. Peptides were desalted using C18 cartridges and dried under vacuum.

Peptides were separated over 60 min using a reverse-phase C18 column (25 cm × 75 μm ID, 1.6 μm, Aurora Series with CaptiveSpray, IonOpticks) on a NanoElute UHPLC (Bruker Daltonics) at 300 nL/min and 50°C. Mobile phase A was 0.1% formic acid in water, and mobile phase B was 0.1% formic acid in acetonitrile. The gradient was set as follows: 2-22% B over 45 min, 22-35% B over 5 min, 35-80% B over 5 min, and held at 80% B for 5 min.

Mass spectrometry was performed on a timsTOF Pro2 (Bruker Daltonics) operating in data-independent acquisition (DIA) mode with Parallel Accumulation-Serial Fragmentation (PASEF). Each acquisition cycle comprised 10 PASEF MS/MS scans, with a capillary voltage of 1400 V and a scan range of 100-1700 m/z. Ion mobility separation was conducted over a 1/K₀ range of 0.7-1.4 Vs cm⁻², with both accumulation and ramp times set at 100 ms. Collision energies were ramped linearly from 59 eV at 1/K₀ = 1.6 Vs/cm^2^ to 20 eV at 1/K₀ = 0.6 Vs/cm^2^. The quadrupole isolation width was set to 2 Th for m/z <700 and 3 Th for m/z >800, using 64 predefined windows spanning m/z 400-1200.

Raw data were analyzed using DIA-NN (v1.8.1) in library-free mode with deep learning-based spectral prediction, employing the Homo sapiens SwissProt database (20,376 entries). Match-between-runs (MBR) was enabled to refine spectral library generation. Peptide and protein identifications were filtered at a false discovery rate (FDR) of <1%. Quantitative protein values were normalized using median normalization. Proteins with missing values in more than one-third of the samples were excluded, and missing data were imputed using a k-nearest neighbors (KNN) algorithm.

**Metabolite Extraction and Full-spectrum Widely Targeted Metabolomics Data** **Processing**

For the extraction of hydrophilic metabolites, approximately 20 mg of ground sample was extracted with 400 μL of methanol/water (7:3, v/v) containing an internal standard. The mixture was vortexed at 2500 rpm for 5 min, left to stand for 15 min, and centrifuged at 12,000 rpm for 10 min at 4°C. The supernatant was collected, stored at −20°C for 30 min, and centrifuged again at 12,000 rpm for 3 min at 4°C. A 200 μL aliquot of the clarified supernatant was subjected to LC-ESI-MS/MS analysis (UPLC: ExionLC AD; MS: QTRAP® System). Chromatographic separation was performed using a Waters ACQUITY UPLC HSS T3 C18 column (1.8 μm, 2.1 × 100 mm) at 40°C with a flow rate of 0.4 mL/min. The injection volume was 2 μL. The mobile phases consisted of water (0.1% formic acid) and acetonitrile (0.1% formic acid) with the following gradient: 95:5 (v/v) at 0 min, 10:90 at 11 min, held until 12 min, returned to 95:5 at 12.1 min, and maintained until 14 min.

For hydrophobic metabolites extraction, samples were extracted with a mixture of methyl tert-butyl ether (MTBE) and methanol (3:1, v/v) containing internal standards. After vortexing for 15 min, 200 μL of water was added, followed by vortexing for 1 min and centrifugation at 12,000 rpm for 10 min at 4°C. A 200 μL aliquot of the upper organic phase was collected and evaporated under vacuum. The residue was reconstituted in 200 μL of acetonitrile/isopropanol (1:1, v/v) for LC-ESI-MS/MS analysis. Separation was carried out on a Thermo Accucore™ C30 column (2.6 μm, 2.1 × 100 mm) at 45°C with a flow rate of 0.35 mL/min. Mobile phases consisted of (A) acetonitrile/water (60:40, v/v) with 0.1% formic acid and 10 mM ammonium formate, and (B) acetonitrile/isopropanol (10:90, v/v) with the same additives. The gradient was programmed as follows: 80:20 at 0 min, 70:30 at 2 min, 40:60 at 4 min, 15:85 at 9 min, 10:90 at 14 min, 5:95 at 15.5 min, maintained until 17.3 min, and re-equilibrated at 80:20 by 20 min.

MS/MS data were acquired using a TripleTOF 6600 system (AB SCIEX) with data-dependent acquisition. During each cycle, the 12 most intense ions (intensity >100) were selected for fragmentation at a collision energy of 30 eV, with an accumulation time of 50 ms per MS/MS event. The electrospray ionization (ESI) source was operated under the following conditions: ion source gas 1 and gas 2 at 50 psi, curtain gas at 25 psi, source temperature at 500°C, and ion spray voltage at +5500 V (positive mode) or −4500 V (negative mode). Additional targeted analysis of hydrophilic and hydrophobic metabolites was performed using a QTRAP LC-MS/MS system (Analyst v2.6.5, Sciex). For hydrophilic compounds, the ESI source parameters were: source temperature 500°C, ion spray voltage +5500 V/−4500 V, ion source gases 1 and 2 at 55 psi and 60 psi, respectively, curtain gas at 25 psi, and high collision gas (CAD). For hydrophobic compounds, the parameters were adjusted to ion source gases 1 and 2 at 45 psi and 55 psi, curtain gas at 35 psi, and medium CAD. Instrument tuning and mass calibration were performed with polypropylene glycol solutions (10 and 100 μM) in QQQ and LIT modes, respectively. MRM (multiple reaction monitoring) was used for data acquisition, with nitrogen as the collision gas (CAD at 5 psi). Specific MRM transitions were scheduled based on the retention time of the corresponding metabolites.

Raw data were processed using MultiQuant software (Sciex) for peak integration and calibration. Peak areas were used to represent metabolite abundances. Metabolites missing in more than one-third of samples were excluded, and missing values were imputed using the KNN method. Quality control (QC) was performed with pooled QC samples containing internal standards to monitor system stability. Metabolite annotation was obtained by integrating the public and self-built databases, including the Human Metabolite Database (HMDB), and the Kyoto Encyclopedia of Genes and Genomes (KEGG) database.

**Supplementary Figures**

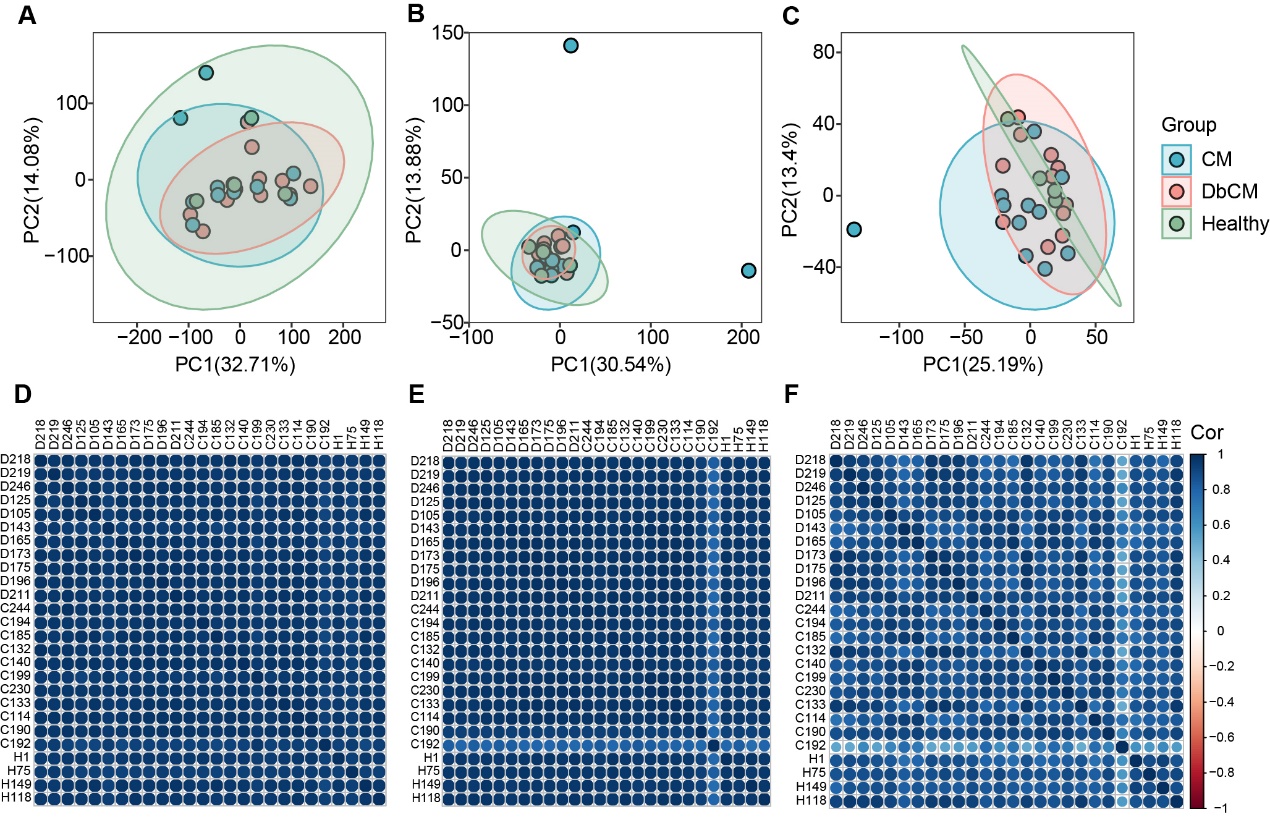

**Supplementary Fig. 1 Overview of multi-omics data from human heart tissues.** (**A-C**) Principal component analysis (PCA) score plots of the transcriptomic, proteomic, and metabolomic profiles, respectively, from cardiomyopathy patients without diabetes (CM group, n=11), diabetic cardiomyopathy patients (DbCM group, n=11), and healthy controls (n=4). One sample from the cardiomyopathy without diabetes group was identified as an outlier in the PCA and excluded from subsequent analyses. (**D-F**) Sample-to-sample correlation matrices for the transcriptomic, proteomic, and metabolomic datasets, respectively.

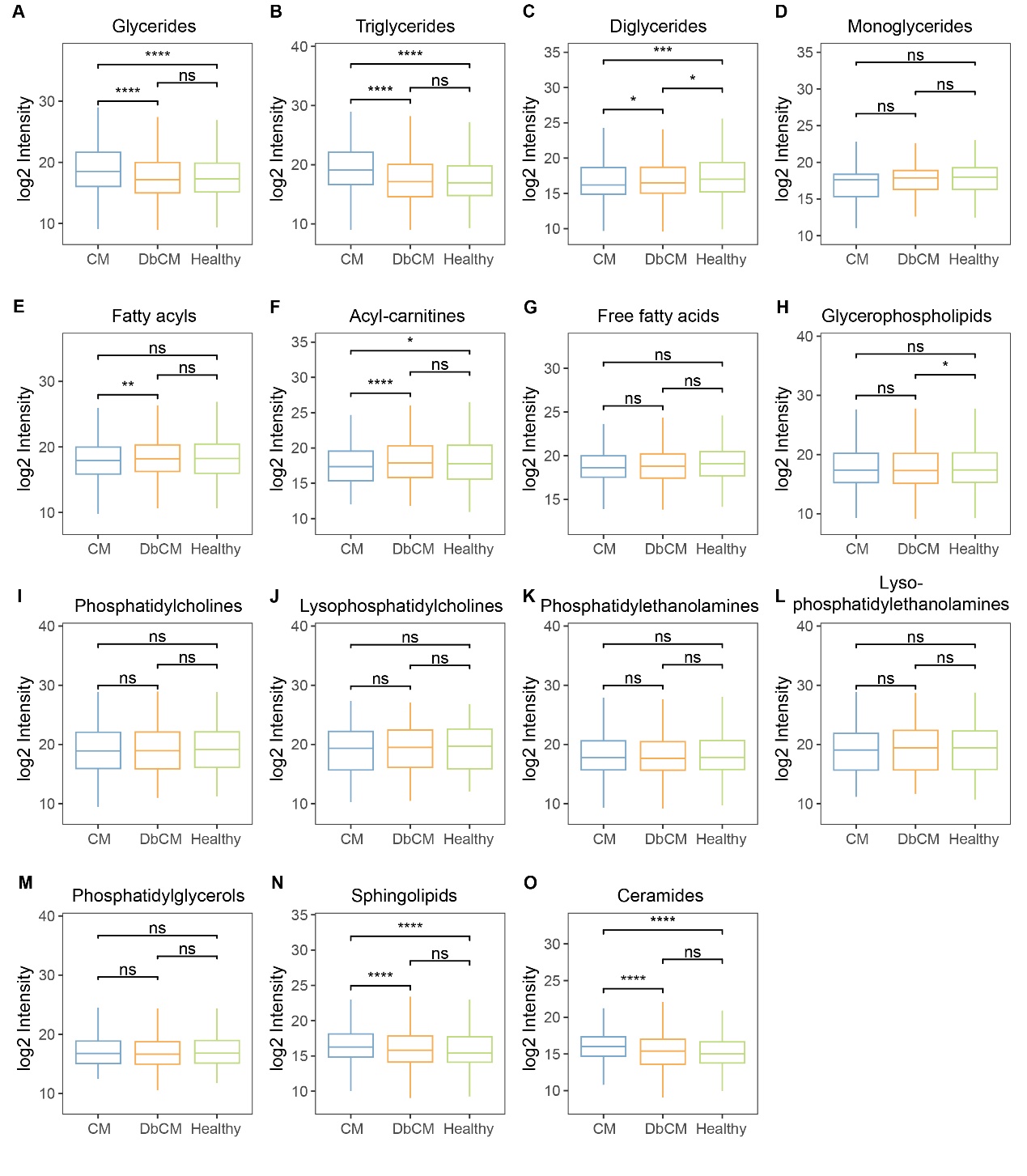

**Supplementary Fig. 2 Box plots of global lipid abundance.** Box plots depicted median values with interquartile ranges (25th to 75th percentiles) of global lipid abundance. Statistical comparisons were performed using two-sided Mann-Whitney U tests. Significance levels were denoted as follows: ns, not significant; **P*<0.05; ***P*<0.01; ****P*<0.001; *****P*<0.0001.

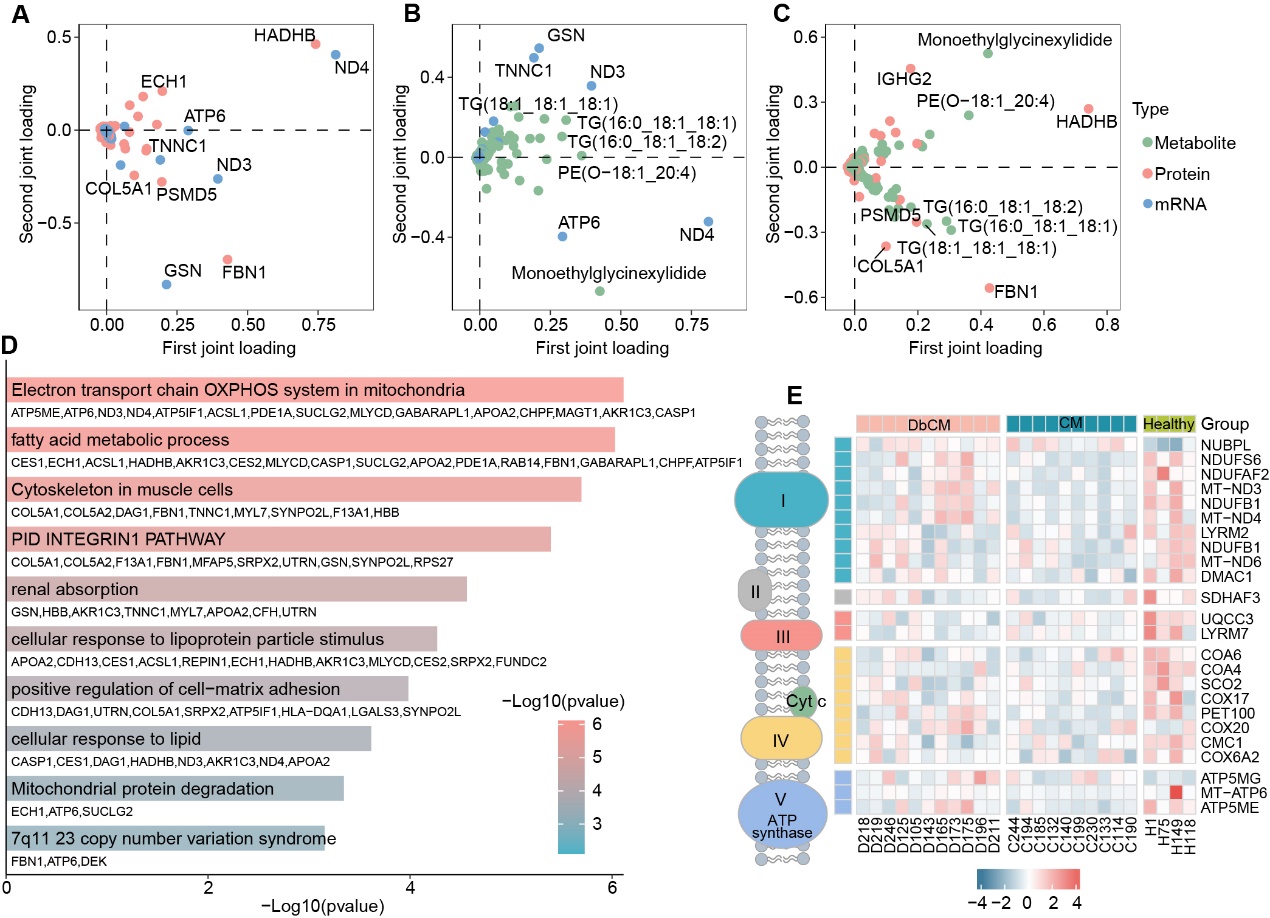

**Supplementary Fig. 3 O2PLS models revealed reveal cross-omics correlations.** (**A-C**) Joint loadings from O2PLS models illustrating correlations between genes and proteins, genes and metabolites, and proteins and metabolites, respectively. Features with the top five squared sums of joint loadings were annotated. (**D**) Pathway enrichment analysis of genes and proteins selected by O2PLS. Dysregulation of oxidative phosphorylation (OXPHOS), fatty acid metabolism, and muscle cell cytoskeleton organization was influenced by multiple omics layers. (**E**) Heatmap of differentially expressed mitochondrial genes (MitoDEGs) and proteins (MitoDEPs) involved in the OXPHOS electron transport chain, showing reduced OXPHOS activity in both disease groups compared to healthy controls. The decrease in OXPHOS levels was less pronounced in the DbCM group.

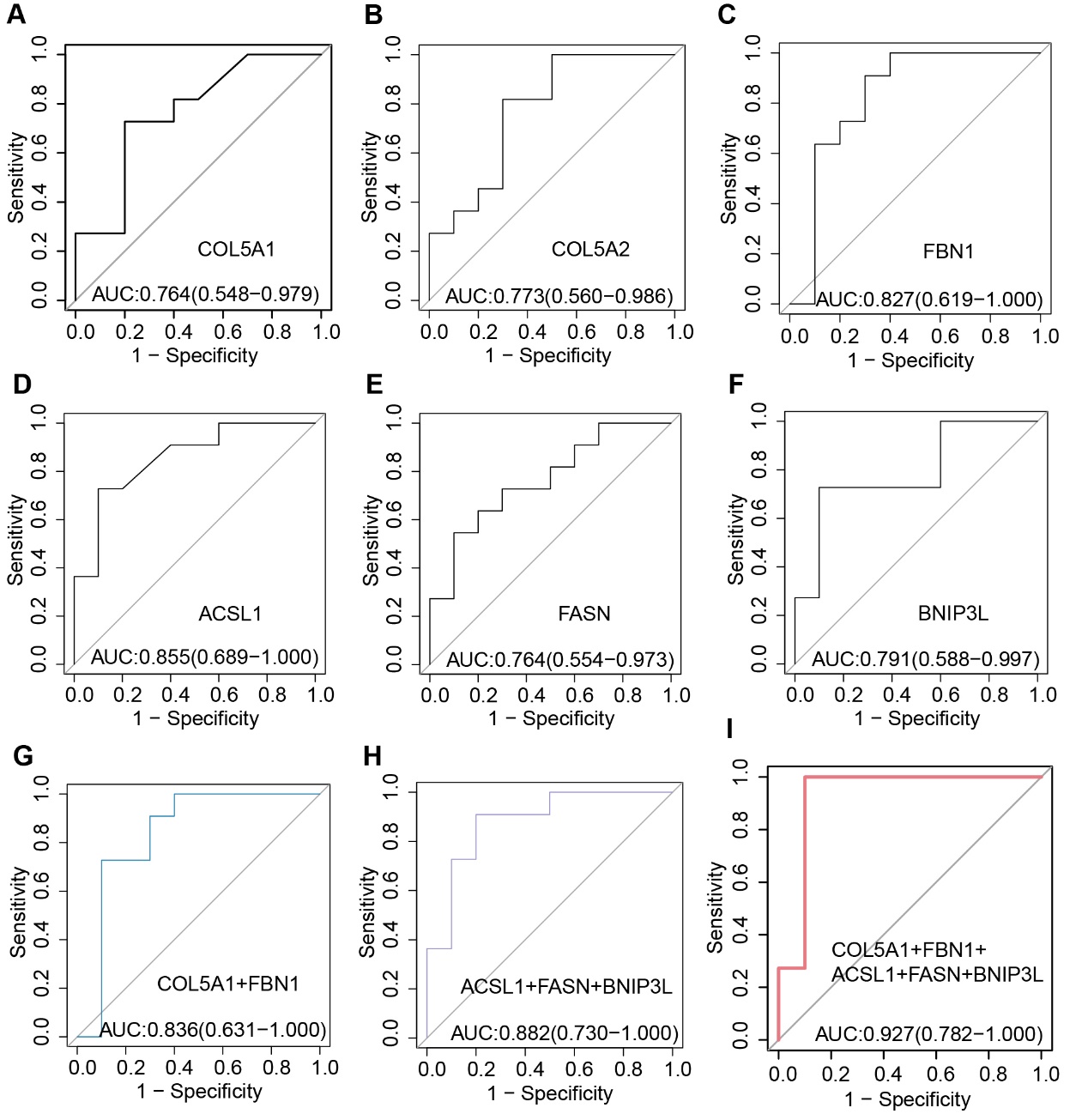

**Supplementary Fig. 4 ROC curves of selected key proteins.** Extracellular matrix-associated proteins, including COL5A1, COL5A2, and FBN1, were evaluated. Due to strong collinearity between COL5A1 and COL5A2 (r=0.953, *P*<0.001), only COL5A1 and FBN1 were retained for the extracellular matrix protein panel based on their biological relevance and literature support. Mitochondrial function-related proteins, including ACSL1, FASN, and BNIP3L, were also selected. A classification model based on these proteins effectively distinguished DbCM patients from cardiomyopathy patients without diabetes.

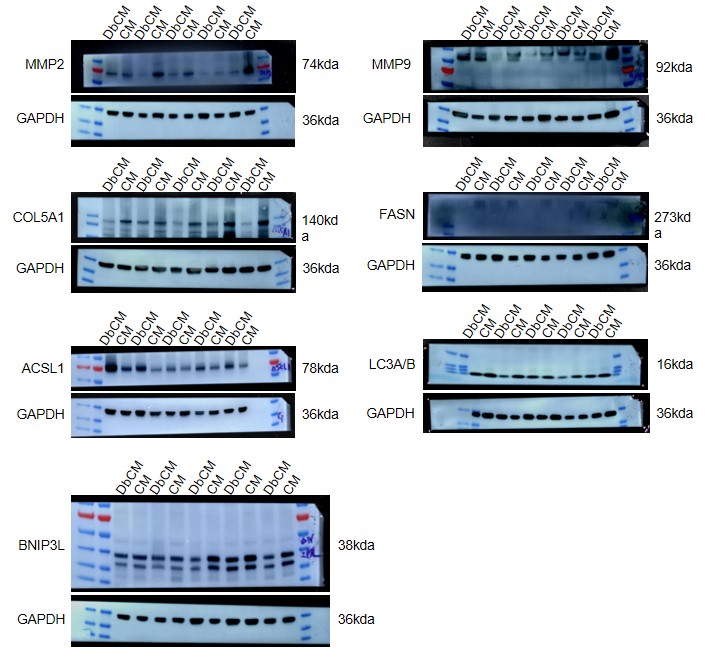

**Supplementary Fig. 5 Unprocessed, uncropped scans of all blots.**
